# Generalised interrelations among mutation rates drive the genomic compliance of Chargaff’s second parity rule

**DOI:** 10.1101/2022.12.23.521832

**Authors:** Patrick Pflughaupt, Aleksandr B. Sahakyan

**Affiliations:** MRC WIMM Centre for Computational Biology, MRC Weatherall Institute of Molecular Medicine, Radcliffe Department of Medicine, University of Oxford, Oxford, OX3 9DS, United Kingdom

## Abstract

Chargaff’s second parity rule (PR-2), where the complementary base and k-mer contents are matching within the same strand of a double stranded DNA (dsDNA), is a phenomenon that invited many explanations. The strict compliance of nearly all nuclear dsDNA to PR-2 implies that the explanation should also be similarly adamant. In this work, we revisited the possibility of mutation rates driving PR-2 compliance. Starting from the assumption-free approach, we constructed kinetic equations for unconstrained simulations. The results were analysed for their PR-2 compliance by employing symbolic regression and machine learning techniques. We arrived to a generalised set of mutation rate interrelations in place in most species that allow for their full PR-2 compliance. Importantly, our constraints explain PR-2 in genomes out of the scope of the prior explanations based on the equilibration under mutation rates with simpler no-strand-bias constraints. We thus reinstate the role of mutation rates in PR-2 through its molecular core, now shown, under our formulation, to be tolerant to previously noted strand biases and incomplete compositional equilibration. We further investigate the time for any genome to reach PR-2, showing that it is generally earlier than the compositional equilibrium, and well within the age of life on Earth.

## 1. INTRODUCTION

In 1950, Erwin Chargaff empirically observed that the four nucleotides are symmetrically abundant across the two strands in a double stranded (ds) DNA molecule – the amount of adenine (A) is equal to the amount of thymine (T), and the amount of guanine (G) is equal to the amount of cytosine (C) – called the first parity rule (PR-1) (1). The explanation of this rule came with the 1953 discovery of the double helical structure of the DNA molecule with the G:C and A:T Watson-Crick base pairings at its core (2). In 1968, Chargaff separated the strands of the *Bacillus subtilis* genome into individual strands and discovered that the same sets of identities found for a double stranded DNA (dsDNA) in PR-1 also holds true on each individual strand of the same molecule, i.e., the amounts of bases are equal in G/C and A/T pairs in each separate strand of a dsDNA as well, formulated as the second parity rule (PR-2) (3,4). In fact, this observation holds true almost universally across extant genomes and has even been extended to the frequency of higher-order oligonucleotides and their reverse complements, where the quantity of each k-mer is equal to the quantity of the reverse complement of that k-mer in the same strand (known as extended PR-2) (5–10). There are, however, exceptions to this rule for most organelles, single stranded DNA (ssDNA) and RNA viruses, which do not comply with PR-2 (7,8,11–13). Being somewhat less clear from the structural and mechanistic considerations of DNA, PR-2 is very robust, hence requires an explanation that is as “crystal clear” and limiting as the Watson-Crick base pairing is for the PR-1. However, reviewing the scientific literature, it seems that an agreed consensus for the exact cause of Chargaff’s second parity rule has not been determined yet. The current hypotheses behind PR-2 can be grouped into two schools of thought: intra-strand symmetry is A) a feature of evolutionary convergence, and B) a feature of the primordial genome. To the best of our knowledge, the sub-categories of explanations are the following: (A) no strand biases for mutation and selection (6,14); selective pressure for the formation of stem-loop structures (15,16); statistical-based approach (9, 17–19); strand inversion/inverted transposition/duplication of dsDNA followed by inversion (13,20); (B) original features of the primordial genome (as opposed to a feature of evolutionary convergence) (21).

Here, we start by re-examining the tolerance to Chargaff’s PR-2 in three kingdoms of species. Reviewing the major explanations proposed so far, we then adopt a completely assumption-free approach to find any possible link between mutation rate constants and PR-2. We demonstrate the contributory role of the equalities of mutation rates in dsDNA that are intrinsically present owing to its complementary double-stranded nature. Showing that, overall, these equalities hold true at a large scale, for at least the human genome, adding a weight on the no-strand-bias (NSB) assumption for PR-2, we also note a substantial variation allowed for the mutation rate constants around the paired equalities. Further examination coupled with symbolic regression and machine learning, leads us to a set of equations that define the universal and more permissive constraints on mutation rates. We demonstrate the better concordance of our equations with the experimental outcomes of genomes both compliant and non-compliant with NSB-driven equilibrium. Furthermore, we elaborate on the evolutionary convergence of genomes to PR-2, explaining why all dsDNA-based life complies with PR-2 at its present snapshot on Earth. The presented work reinstates the mutation rates as the major drivers behind the emergence of PR-2, demonstrates the simple principles behind the complex question, and can serve as an important basis for the future molecular evolution studies in genomics.

## MATERIALS AND METHODS

### General notes on the performed calculations

The developed workflows and analyses in this study employed the R programming language (22). The resource demanding computations were performed on a local Linux-based computing cluster at MRC WIMM, University of Oxford, by using nodes with 3 × 2.7 GHz 8-core E5-2680 Intel Xeon processors and 256 GB random access memory. The analytical derivations and checks have been done *via* the Mathematica software (44). The server application (ATGC Dynamics Solver) was written in R, using the Shiny library (23) and server application (http://shiny.rstudio.com). Figures were created with the R base (22), ggplot2 (24) and gridExtra (25) libraries. Handling of the datasets were done by using the R base, tidyverse (26) and reshape2 (27) libraries.

### Processing of the eukaryotic, prokaryotic and DNA virus species

We have examined over 8000 genome sequences ranging from eukaryotes, prokaryotes, and DNA viruses. We accessed the Ensembl Genomes database *via* FTP (ftp://ftp.ensemblgenomes.org) and downloaded all genome sequences from, at the time of performing this work, the latest release for bacteria (ftp://ftp.ensemblgenomes.org/pub/bacteria/release-48) and eukaryotes (ftp://ftp.ensembl.org/pub/release-102/fasta/). For DNA viruses, we accessed the NCBI database (https://www.ncbi.nlm.nih.gov/labs/virus/vssi/#/find-data/virus) and downloaded all species categorised as “DNA viruses”. For the bacteria and eukaryotes, we downloaded the genome sequences in an iterative process *via* the R code. We note that a significant number of sequences had to be removed because of, for instance, duplicates of a given strain of a given species. Taking prokaryotes as an example, there were approximately 44,000 species in the original file from the Ensembl database, but we noticed that a single species could have thousands of entries in the file which would skew the species-wide analysis and yield inaccurate results. The same problem also occurred with eukaryotes and DNA viruses. Thus, the goal was to filter the files with the following methodology: in a group of species with the same name, we only saved the first occurring row of the given species name and repeat this process for the remainder. At the end of this process, we discarded many duplicates and filtered the data as consistently as we could, including manual filtering processes. Full details are on the corresponding GitHub repository.

### Processing of the human and Chimpanzee genomes

Analyses involving the human genome were performed using the unmasked version of the human reference sequence hg19/GRCh37, as accessed from Ensembl (28) genome database (www.ensembl.org). We used the unmasked chimpanzee genome, version 2.1.4.75, for calculating the genomic base and dyad contents (**Supplementary Note S1**). The fasta files were downloaded from the Ensembl database. While also trying the masked genome, we noted only negligible differences on both single base and dyad contents. For counting dyads, both segmentation (into dimers) and sliding window methods were used, again with no significant differences noted.

### Mutation rate constants for the use in simulations

The mutation rate constants used in this work were all brought from previous work using the Trek methodology (29), where the human genome mutation rates were revealed through the remnants of LINE-1 elements in a single genome manner. We also re-calculated such rate constants using LINE-1 elements that reside in only +, and in only –, strands to arrive to strand-specific but still genome-wide overall values for mutation rates. Strand-symmetry accounted mutation frequencies for the following species: *Homo sapiens*, *Escherichia coli*, *Caenorhabditis elegans*, *Drosophila melanogaster, Aotus thaliana*, and *Saccharomyces cerevisiae* were obtained from (30). All such frequencies were next brought into a scale of our rate constants for the use in our simulation models. In the supplementary materials of (29), **Figure S3** shows the relationship of the mutation rate constants for the human genome obtained with the Trek methodology versus the mutation frequencies reported from other datasets. We replicated **Figure S3a** (29) with the obtained frequencies from other species and the mutation rate constants from the Trek methodology, and performed a linear regression through the zero origin. This resulted into the *k* = 2.831*f* equation. As we were interested in the genomic average mutation rate constants in a time domain, we used an assumption that this quantitative relationship in between the mutation rate constants and mutation frequencies holds true for the other (non-human) five species. Thus, the frequency values, obtained from the literature, were used as inputs *(f)* in the above equation to generate the average mutation rate constants (*k*) in time domain for our simulation.

### Generation of simulations

The numerical analyses were performed using the R programming language with its deSolve library for solving the system of differential equations. Random sampling of the mutation rate constants for the continuous uniform distribution were done with the runif() function, and the truncated normal distribution were done with the rtruncnorm() function from the truncnorm library (31). All simulations were run in parallel execution using the foreach() (32) and doParallel() (33) functions. For reproducibility, we set the initial seed to 1 and used the doRNG (34) library for the parallel child processes of in our foreach loops.

### Defining the equilibration time periods

Simulation of the replicates were done using symmetry-constrained mutation rate constants drawn from a truncated normal distribution using the rtruncnorm() function from the truncnorm library in R. Where the equilibrium was needed, we carried out the following five-step process to define a genome equilibration tolerance for the simulation: A) calculate the sum of the differences of base contents divided by four; B) repeat the simulation 100,000 times; C) calculate the mean fluctuation of (A) for the 100,000 times; D) check that the mean fluctuation is not greater than 1% of 25 of the mean fluctuations that we define as the EQtolerance; E) calculate the absolute difference of each base content and find the difference between the maximum and minimum. When this time is less than or equal to the EQtolerance, genome was considered equilibrated. Independently from the equilibration compliance, PR-2 compliance was defined as the time when both the GC skew and AT skew are below their respective tolerance values obtained for each of the three kingdoms: eukaryotes, prokaryotes, and DNA viruses, as discussed below.

## 2. RESULTS AND DISCUSSION

### 2.1. The universality of PR-2 in dsDNA genomes of different species

The re-examination of the universal nature of the Chargaff’s PR-2 compliance of dsDNA on an up-to-date collection of genomes is shown in **Figure 1a**, where we bring the plots for *C_GC_* vs. *C_AT_* skews, as defined by (G-C)/(G+C) and (A-T)/(A+T), for the genomes of species across the eukaryotes, prokaryotes, and DNA viruses. A strong adherence to PR-2 can be noted in all kingdoms (**Figure 1a**) with the intra-strand symmetry phenomenon holding true across the extant dsDNA genomes (standard deviations: 1.908×10^−3^, 1.907×10^−2^, 9.666×10^−2^ for *C_GC_*, and 1.178×10^−3^, 9.400×10^−3^, 1.041×10^−1^ for *C_AT_* across eukaryotes, prokaryotes and DNA viruses, respectively), except for the organelle and other ssDNA genomes (7–10). While focusing on the low-density scatter in the *C_GC_* and *C_AT_* skews, the viral DNA genomes are narrowly spread, while the eukaryotic organisms form a dense and localised cluster. In prokaryotes, the low-density scatter in the plots is not only more diffuse, as compared to eukaryotes, but there seems to be an emerging diagonal pattern, linked, based upon our experimentations, with the G+C content of a genome.

**Figure 1.**
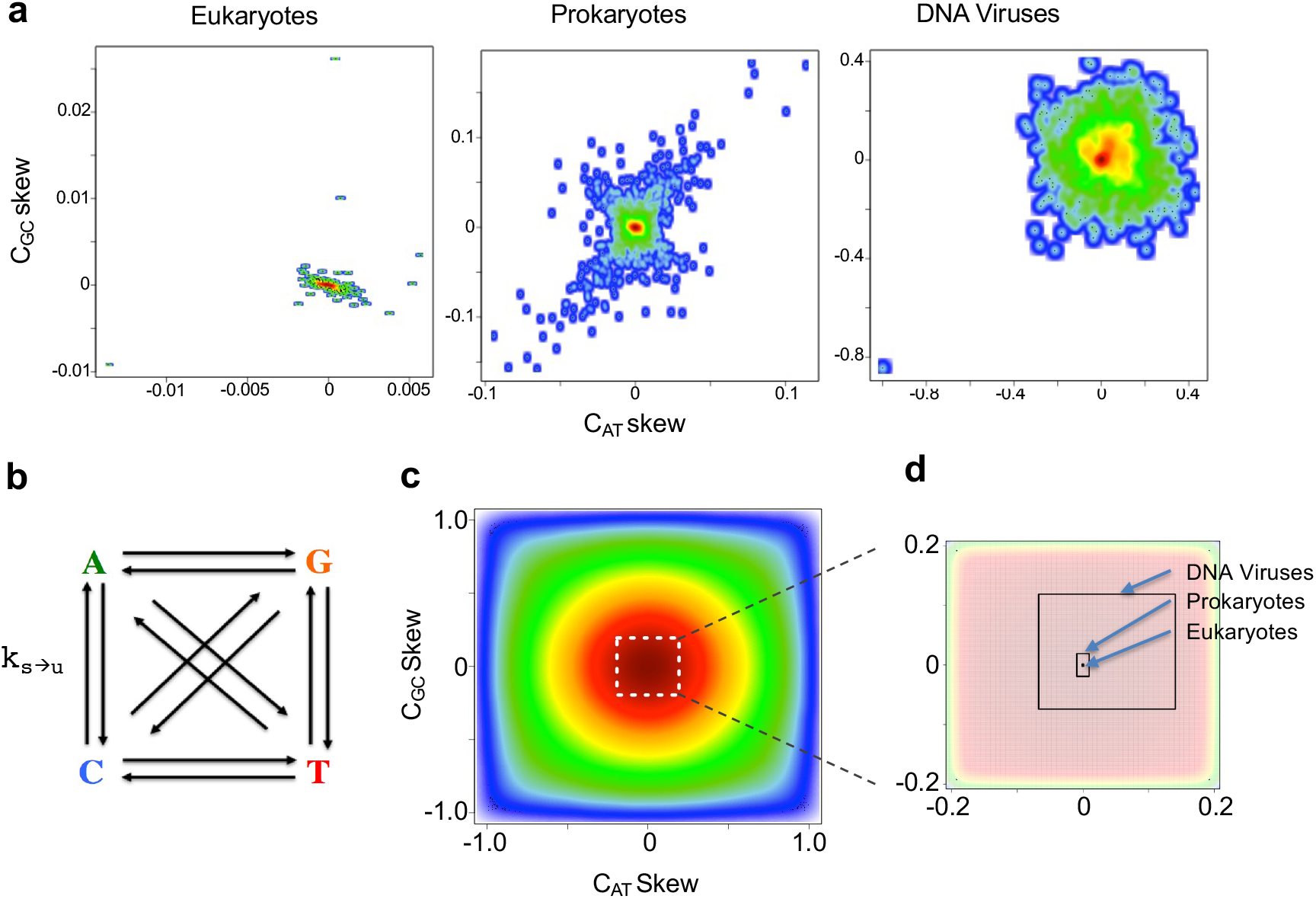
Chargaff’s PR-2 in different species and the unconstrained mutational network model. (a) The plots for the base content GC vs. AT skews for each of the three kingdoms analysed in this work (see **Figure S2** for the exact mean and standard deviation values). The 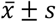 range for each kingdom was used for the naturally observed PR-2 fluctuation, while analysing the results of the simulations. (b) The most general mutational network model, where all the 12 *k*_*s*→*u*_ rate constants are unique and independent from each other. The system was numerically solved to produce the time evolution of genomic base content within 4.28 byr period, the current maximum estimate of age of life on Earth (43). (**c**) The 2-dimensional kernel density estimate scatterplot, showing the distribution of GC and AT skews from the outcome of the simulation (colours vary with decreasing occurrence frequency from red to blue). There, the white dotted box indicates the area zoomed in **d**. (**d**) Zooming to demonstrate the strict PR-2 compliant zones for each of the three kingdoms, where dimensions of the boxes represent the total 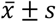 range for the corresponding axis and kingdom (see **Section 2.1** for exact values).

Based on these plots, we can infer the allowed PR-2 fluctuations for each of the three kingdoms. This allows the definition of “tolerance” values for PR-2 compliance, essential for our further studies. We examined the tolerance values from the *C_GC_* and *C_AT_* skews as opposed to the *C_G/C_ and C_A/T_* ratios (see **Figure S2**), because, in the former case, the range of allowed values are limited between [0, 1] while the mathematical nature of the *C_G/C_ and C_A/T_* ratios allow a biased range from [0, ∞]. The preferential regions of PR-2 compliance were defined from the analysed experimental genomes for each kingdom, by taking the mean value of the skews (usually tending to 0) and one standard deviation allowed at both sides of the mean. For *C_GC_ skew*s, the PR2-compliant preferential fluctuation regions were 7.780×10^−5^ ± 1.908×10^−3^, 1.425×10^−4^ ± 1.907×10^−2^, and 2.262×10^−2^ ± 9.666×10^−2^ for eukaryotes, prokaryotes and DNA viruses respectively. For *C_AT_ skew*s, the ranges were 1.096×10^−5^ ± 1.178×10^−3^, - 1.386×10^−5^ ± 9.401×10^−3^, and 3.668×10^−2^ ± 1.041×10^−1^ in the same order as above.

### 2.2. An overview of the past theories to explain PR-2

One of the earliest attempts to explain the emergence of PR-2, and one that was initially well received, was the hypothesis of no-strand-bias (NSB) conditions (6,14). In 1995, Sueoka determined that, when there is no bias in mutation and selection between the complementary strands, base mutation may explain the parity phenomenon (14). For instance, for a *C* → *T* mutation in one strand of a dsDNA under NSB, the rate for the *C* → *T* mutation on the other, complementary, strand can be considered the same. Due to the first parity rule, C➔T mutation leads to a *G* → *A* mutation on the complementary strand. Therefore, the *C* → *T* and *G* → *A* mutation rates match on the same strand too, since *G* → *A* mutation is the same *C* → *T* mutation on the complementary strand, and the rates of *C* → *T* mutations on both strands match under NSB. As a result, a mutation network of 12 unique mutation rates can be simplified down to only 6 independent rates (14). In the same year, Lobry derived a set of differential equations that showed that, at equilibrium, the base frequencies within each strand are equal, i.e., resulting in PR-2 (6). While a mutation network with NSB was a working hypothesis by both Sueoka and Lobry, it was shortly afterwards rejected by Lobry because the equations assume that both the mutation rates and the resultant genomes are constantly at equilibrium. It suggests that any deviation from PR-2 could either mean that the model is wrong or that the system is out of equilibrium. Considering known examples of genomes being out of equilibrium (35–37), as well as mutational processes bearing some strand asymmetries (38), this hypothesis cannot be the universal explanation for PR-2 (36,37). While Sueoka and Lobry used a base-mutation model to explain the cause of PR-2, in the same year, Forsdyke proposed that PR-2 is a product of selective pressure favouring point mutations that assist in the formation of stem-loops (15,16). He argued that such formations may be advantageous for recombination events, thus playing an important role in driving the symmetry between complementary oligonucleotides. However, similar to Sueoka and Lobry, Forsdyke’s hypothesis has also been challenged. Zhang et al. suggested that most oligonucleotides (with sequence length of five and longer) do not have a reverse complement in close proximity in the genome sequence and thus the short-range contribution of local stem-loop potential by complementary oligonucleotides is limited (39). Chen et al. also challenged the stem-loop hypothesis for human chromosomes (40).

Albrecht-Buehler proposed that PR-2 is the inevitable, asymptotic product of the cumulative action of proposedly ever-repeated inversions and inverted transpositions in genomes (20). He extended his inversion/transposition hypothesis by observing that these mechanisms also generated a universal triplet profile for most of the evolving organisms - i.e., the frequency of triplet oligonucleotides are almost equal to the frequency of their reverse-complement, in compliance to the extended PR-2. He further explained that the evolutionary advantage of an almost universal genome may be to help with horizontal gene transfer between species that are vastly different from one another. His conclusions from this paper, however, have been challenged by Zhang et al. who found that, at least among prokaryotic genomes, two common triplet profiles existed, one for low-G+C and the other for high-G+C content genomes (41). Similar to Albrecht-Buehler’s hypothesis, Okamura et al. proposed that inversions, which may or may not be preceded by duplications, may be a major contributor to the phenomenon of the intra-strand parity rule (13). They found that when subjecting the human mitochondrial DNA to inversions *in silico*, the frequencies of specific trinucleotides and their reverse complementary oligonucleotides became broadly symmetrical, suggesting that this mechanism could lead a sequence to parity. It is worthy to note, however, that the biological and mechanistic plausibility of such repeated inversions and transpositions as a widespread phenomenon in genomes remains to be confirmed.

A number of scientists have used statistics-based approaches to explain the cause of PR-2. For instance, Baisnée et al. (7) demonstrated that using simple, first-order Markov models to model biological sequences was not enough to explain symmetries in dsDNA genomes. Instead, the emergence of symmetry may be explained through a combination of mechanisms, such as inversions, among other possibilities. Hart et al. (17) showed that the Gibbs distribution, which is associated with the reverse complementary relationship between the nucleotide interactions on each strand of dsDNA, can explain PR-2. An observation made by Shporer et al. (8) showed that when we consider a genome of a given length *L*, then the highest k-mer (substrings of length *k*) for which PR-2 holds true is given by 0.7ln(*L*), thereby demonstrating that the extended PR-2 rule holds true up to a certain length of a k-mer. However, their observation has been generalised in a recent paper by Fariselli et al. (9), which does not impose an upper limit to the k-mer length for the PR-2 to hold true. They proposed a solution based on a maximum entropy approach, which predicts that at equilibrium, in a long-enough dsDNA sequence, the probability of occurrence of a k-mer and its reverse complement tend to be the same. In other words, the leading force shaping the symmetrical DNA duplex structure is randomness, rather than biological/environmental pressures. A recent paper by Cristadoro et al. (10) showed that Chargaff’s parity rules are not the only symmetry phenomena present in the genetic sequences of *Homo sapiens*, instead, there exists a hierarchy of symmetries at different structural scales. In fact, these observations are similar across all nuclear chromosomes of *Homo sapiens*, thus suggesting that some mechanisms that shape both the structure and symmetry work at the same time in the chromosomes, the leading mechanism of which is currently unknown. They further investigated the evolution of the genome dynamics, in which they developed a model to mimic the action of inversion/transpositions of transposable elements on DNA, originally motivated by Albrecht-Buehler’s simulations (20). The results show that the simultaneous occurrence of symmetry and structure is an emergent property of the dynamics of transposable elements in DNA sequences (9). In particular, they found that symmetry and structure change differently depending on the time scales, i.e., for a large time interval, they were able to reproduce the same structure and symmetry as in extant genome sequences.

An alternative school of thought on the cause of PR-2 was proposed by Zhang et al (21). They suggest that the origin of strand symmetry and oligonucleotide frequency conservation has existed from the very beginning of the genome evolution and thus, compositional features of the genome would be “relics” of the primordial genome as opposed to a feature of evolutionary convergence - the umbrella hypothesis of the previously discussed scientific literature. Zhang et al. argued that the primordial genome would be composed of approximately equal amounts of uniformly distributed forward and their reverse-repeated sequences, thus resulting in a strand symmetric genome where frequency conservation is a consequence of it, although they note that the degree of strand symmetry decreases with increasing order of oligonucleotides.

### 2.3. Assumption-free approach to look for a link between mutation rates and PR-2

Of all the prior hypothesis reviewed above, we can see that the NSB approach (6,14) was the one that was proposing a strong constraint at the core of the molecular processes that govern dsDNA composition, thus being as strict as the observation of the PR-2 compliance in genomes is. However, the approach was dismissed (36,37) due to the expected aberrations from NSB (38,42) and/or compositional equilibrium (35) regionally, in shorter dsDNA spans, and in some species. In this work, we thus revisit the link between the mutation rate constants and PR-2, but with no initial assumption on the mutation rates and their equalities whatsoever. We numerically calculated the base content coming out of a 4.28 billion years (byr, the estimated age of life on Earth (43)) dynamics of unconstrained cross-mutational network, where every base mutation is assumed to have an independent rate constant, i.e. each conversion arrow in **Figure 1b** to have an unconstrained k_s➔u_ value, where the subscript denotes the mutation of the base *s* into *u*.

We simulated 25 million systems starting from a completely random genome (0.25 for {*A*, *T*, *G*, *C*} base compositions) to obtain enough PR-2 compliant samples based on our PR-2 tolerance ranges calculated in **Section 2.1** for the eukaryotes, prokaryotes, and DNA viruses. We ran the simulation four times separately by randomly drawing the mutation rate constants from a uniform distribution. To allow for a broad range for our random rate constants, we took the value of the symmetry-corrected maximum mutation rate (*k*_*C*→*T*_/*k*_*G*→*A*_) observed in human genome from (29), added its standard deviation additionally scaled by multipliers 1, 2, 5, and 10. This resulted in [0, 1.799], [0, 2.490], [0, 4.561], and [0, 8.013] ranges, for all the multipliers respectively. The discussion below operates on the outcomes with the multiplier 1, but we verified that the results did not differ much for the other tried cases. We did this for the following reason: while simulating the cases and allowing variation in the mutation rate constants, one approach would be to take the human rate constant values and fluctuate around them. However, as we do not know what kind of fluctuation is relevant in the original life, it is better to simulate cases where we go beyond the certain range, hence scaling the random sampling of the mutation rate constants.

In this most general case, all 12 rate constants are unique, and the corresponding system is numerically evolved using the following four kinetic equations:

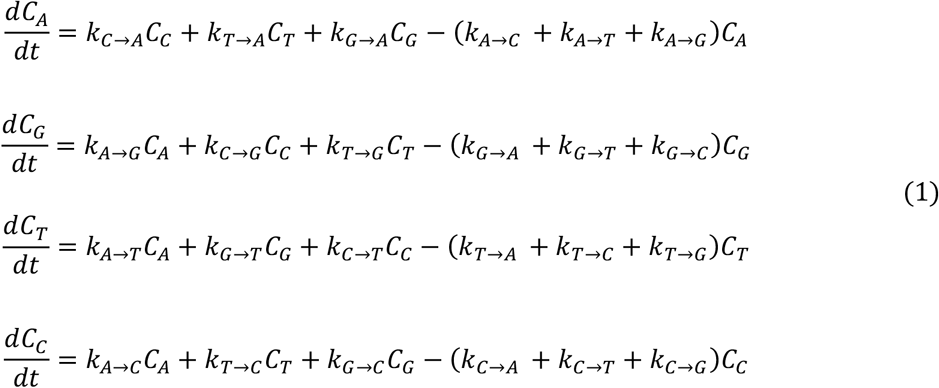

where *C_s_* is the content of the base *S*, and *k*_*s*→*u*_ is the mutation rate constant for the *s* → *u* mutation. Recording the base contents at the end of the 4.28 byr period, we can see that the final *C_GC_* and *C_AT_* skews have a wide range of distribution, peaking at 0 (**Figure 1c**). That preference towards 0 is, however, far from the observed PR-2 compliance in actual genomes, where the compliance region is much narrower at around the 0 value for the skews. This is demonstrated in **Figure 1d**, where the centre of the plot is zoomed with the compliance regions highlighted for eukaryotes, prokaryotes and DNA viruses.

Two other scenarios were also tried for the simulations, where every independent mutation rate constant is assumed to have A) a normal distribution centred at their corresponding average values for *Homo sapiens*, and B) symmetry-constrained rates from a normal distribution centred at symmetry-averaged rates for *Homo sapiens*. The values for the mutation rates were taken from (29) and summarised in **Table 1**. For each of the four separate simulations, the standard deviation values were scaled by *m* ∈ {1, 2, 5, 10} multiplier values to examine the various spread scenarios around the averages.

**Table 1.** Rate constants used in the simulations for i➔j mutations. There, the notation such as AC denotes a rate for the *A* → *T* mutation. The Individual singleton rate constants used in simulation (A) are brought in the upper part. Those are followed by the strand symmetry-averaged singleton rates used in simulation (B). Mutation rate constants were taken from (29).

| Mutation | Mean | SD |
| --- | --- | --- |
| AC | 0.198 | 0.100 |
| TG | 0.199 | 0.098 |
| AG | 0.679 | 0.553 |
| TC | 0.875 | 0.824 |
| AT | 0.210 | 0.111 |
| TA | 0.216 | 0.132 |
| CA | 0.371 | 0.505 |
| GT | 0.269 | 0.139 |
| CG | 0.304 | 0.158 |
| GC | 0.238 | 0.117 |
| CT | 1.173 | 0.894 |
| GA | 1.044 | 0.387 |
| AC\|TG | 0.198 | 0.099 |
| AG\|TC | 0.753 | 0.673 |
| AT\|TA | 0.212 | 0.119 |
| CA\|GT | 0.322 | 0.378 |
| CG\|GC | 0.273 | 0.144 |
| CT\|GA | 1.109 | 0.690 |

These results are summarised in **Figures S3**-**S4**. We note that, at 4.28 byr time point, the solutions to the simulation with the symmetry-constrained rates (B) exhibit a distinct difference in behaviour, as compared to the simulations in (A). The simulation (B) reveals preferential islands in *C_GC_ vs C_AT_ skew* plots (**Figure S4**), and a diagonal spread that is scaled as we increase the standard deviation values for the mutation rate sampling range. Interestingly, the diagonal pattern is found in the actual skew plot for prokaryotic organisms (**Figure 1a**).

### 2.4. *De novo* link between the unconstrained mutation rates and PR-2

For the 25 million *in silico* generated systems (**Section 2.3**), we obtained the combinations of rate constants that resulted in PR-2 compliant base contents after 4.28 byr. Examining the *C_GC_* and *C_AT_ skew*s at the end of the simulations (**Figure 1c**), we can zoom onto the area that is comparable to the selected PR-2 compliance regions, as inferred from the experimental genomes (**Figure 1a**) defined in **Section 2.1**. **Figure 2** shows the ratios of *k*_*s*→*u*_ pairs that are expected to be equal under the NSB assumption, but not pre-set as equal in these simulations. We show the distributions of the ratios of *k*_*s*→*u*_ pairs for all the generated combinations of rate constants in **Figure 2a** that end up with the system complying with PR-2. Those in **Figure 2b** show a classical ratio distribution shape heralding the division of two uniform distributions (**Figure 2b**). Zooming onto the strict PR-2 compliance region, and investigating the rate constants that resulted in low *C_GC_* and *C_AT_ skew*s in our simulations, the above-defined rate constant ratios become distributed differently, peaking at the value of 1, though still showing a significant spread (**Figure 2a**). Hence, with no constraint on any interdependence amongst the mutation rates (inter-mutation network brought in **Figure 1b**), we can see that the combinations of those 12 independent rate constants that allow for a PR-2 compliance reflect, at least in part, the NSB rate constant equalities (**Eqs. 1**). Importantly, the combinations still allow a significant variation at around those equalities (**Figure 2a**), potentially governed by higher degree of interdependences in mutation rates that we investigated in further sections.

**Figure 2.**
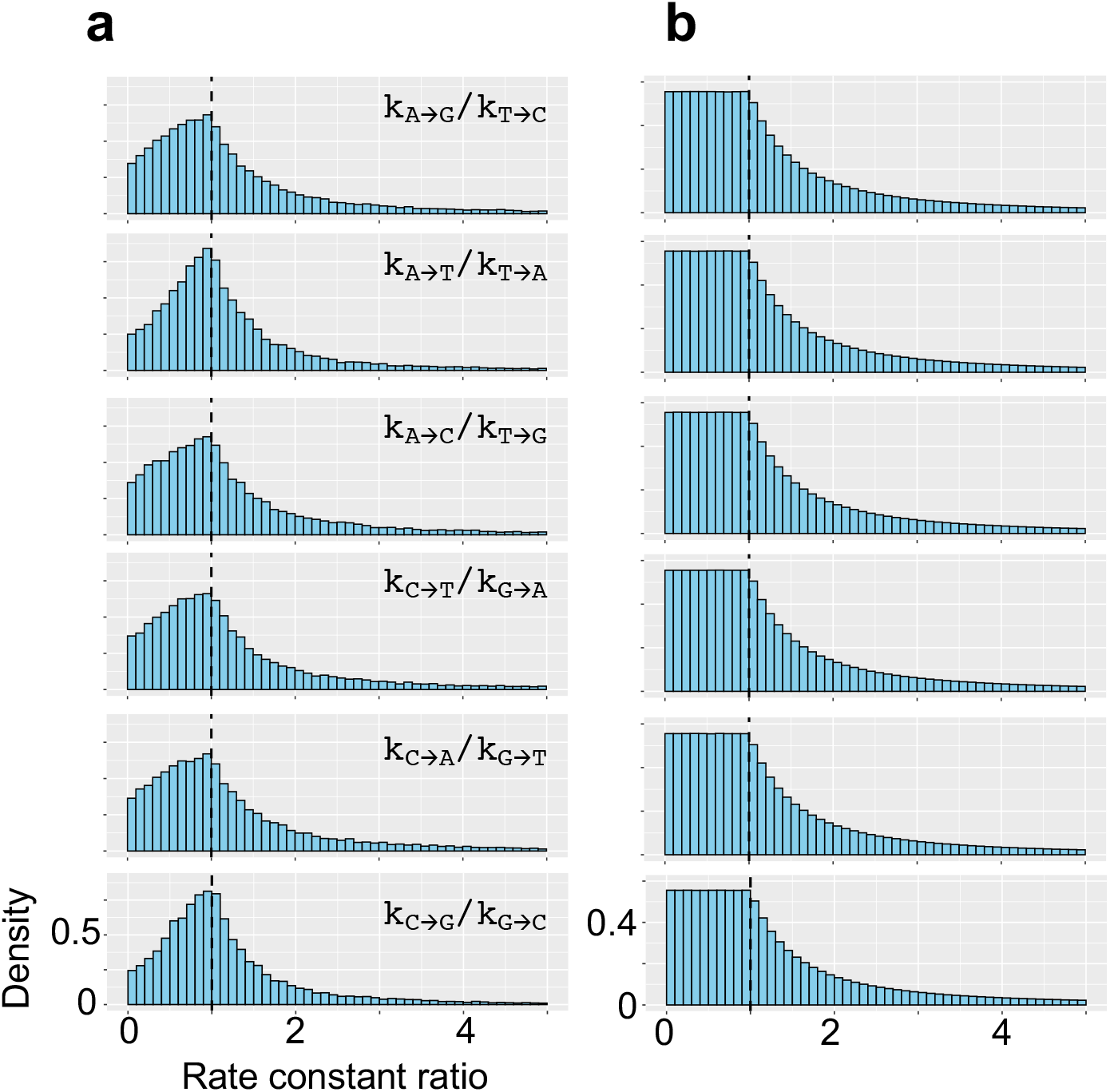
Distributions of the rate constant ratios from the numerical analysis of the unconstrained mutational network model. This represents the most general model, where all the 12 *k*_*s*→*u*_ rate constants (**Figure 1a**) are unique. The system is numerically solved to produce the time evolution of genomic base content within 4.28 byr period, the current maximum estimate of age of life on Earth (43). The plots **a** and **b** show the distributions of the ratios of *k*_*s*→*u*_ pairs that are expected to be equal (**Eqs. 1**) according to the no-strand-bias (NSB) assumption, but not pre-set as equal in the simulations. The set of histograms in **a** show such ratios for all the rate combinations in the PR-2 compliance zone peak at the value of 1, showing that the solutions tend to the compliance with the parity rule when the rate constants tend to satisfy the equalities stemming from the NSB assumption, though also showing a substantial variation. The set of histograms in **b** show such ratios for all the rate combinations except those in **a**, the compliance zone, and have the classical ratio distribution shape, which can be obtained by dividing two uniform distributions.

### 2.5. Link between the mutation rates and PR-2 under the no-strand-bias assumption

From the **Section 2.4** and **Figure 2a**, we inferred that there is a wide range of combinations in mutation rates that can result in genomes that exert Chargaff’s PR-2 as a phenomenon emergent from those rate constants within the timeframe of the age of life on Earth. However, a part of the PR-2 compliant rate constant combinations, springing from our unconstrained simulations, showed rate equalities similar to NSB assumption. Therefore, before going on to investigate the more generalised rules that may exist to link mutation rates with PR-2 compliance, here we first outline the simpler model under NSB assumption. As discussed in prior literature, equalities of certain mutation rates could be present in DNA owing to its complementary double-stranded nature, if we assume NSB for mutation rates (6,14). To clarify how the mutation rate equalities emerge, consider the example of *k*_*C*→*T*_ = *k*_*G*→*A*_ (**Figure 3a**). Central to the NSB model is a plausible assumption of the strand-invariance of the mutation rates, i.e. the *C* → *T* mutation happens at the same rate independently from whether *C* is in the template or complementary strand of the dsDNA. Therefore, each strand will also have the complementary *G* to *A* conversions with the rate similar to *C* to *T* conversion, hence *k*_*C*→*T*_ = *k*_*G*→*A*_. This symmetry in rate constants significantly simplifies the mutation network from 12 to 6 independent rate constants:

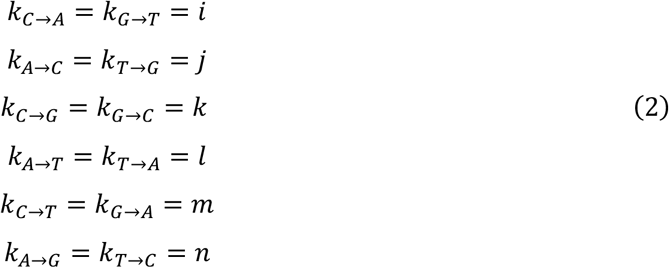

**Figure 3.**
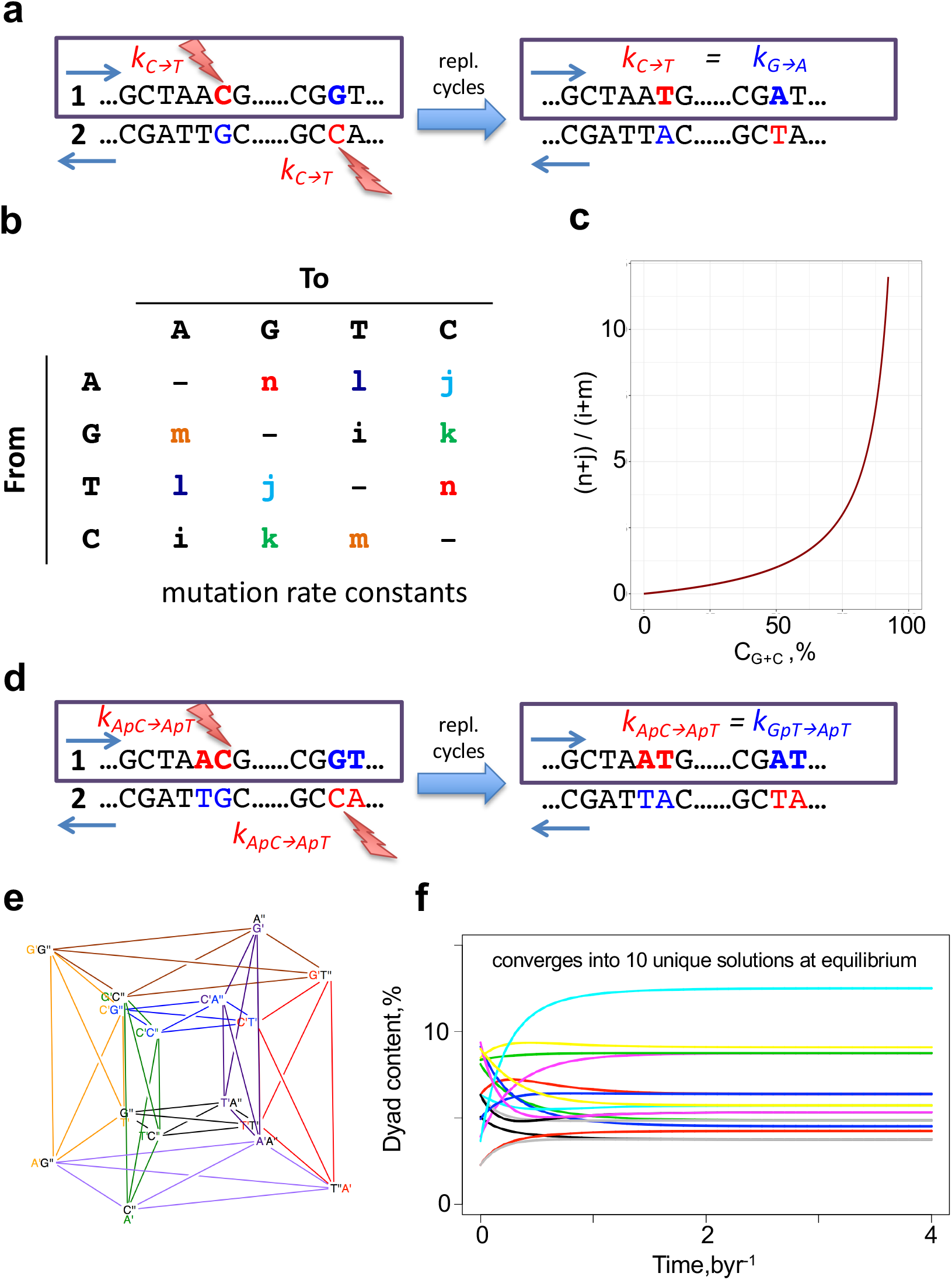
The no-strand-bias (NSB) assumption for singletons and dyads. (**a**) Illustrates the NSB principle, on the example of *C* → *T* transition. (**b**) The mutation rate matrix with only 6 independent parameters. The inferred dependency between the genomic equilibrium G+C base content and mutation rate constants is shown in **c**. (**d**) Example of the NSB model in a dyad context. In the oligomeric context, NSB happens where both the source and the substituted states of two mutations are reverse complementary to each other. The figure demonstrates this, analogous to the single-base case discussed before, on the example of the equalities between the rates of *ApC* → *ApT* and *GpT* → *ApT* dyad mutation rates. (**e**) The schematic representation of the cross-mutation network among dyads is constructed based on the tesseract (hypercube). There, the primed superscripts denote the position in the dyad, from 5’ to 3’ direction (A’G” is the same as ApG). Each line should be interpreted as a set of two counter-directed arrows. (**f**) shows an example of the time evolution of dyad contents starting from an arbitrary set of initial dyad contents and rate constant values. The colouring scheme is random.

The reduced number of independent rate constants from the NSB assumption produces a simpler mutation matrix (**Figure 3b**), and a simpler system of kinetic equations:

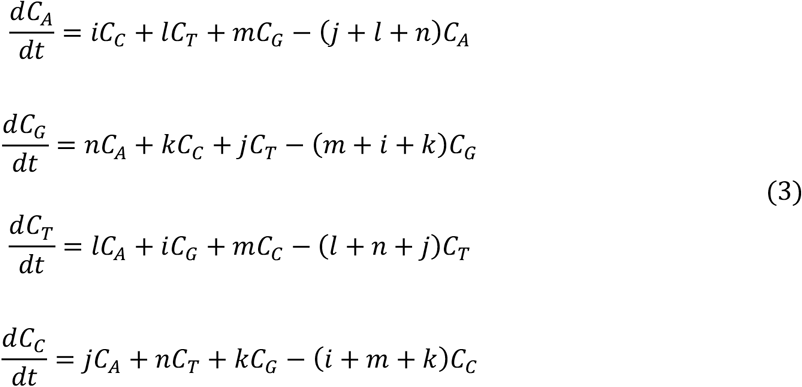

each describing the evolution of A, G, T and C base contents in fractions. At equilibrium, the system is fully solvable, and the equilibrium base contents are given through the following equations (see **Supplementary Note S1** for the derivation):

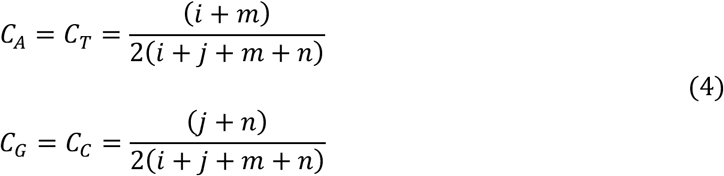

The solutions imply the *C*_*A*_ = *C*_*T*_ and *C*_*G*_ = *C*_*C*_ equalities for base composition at equilibrium under NSB for mutation rates. These solutions also link the mutation rate constants (a molecular level characteristics of a DNA) with the equilibrium genome composition. We can express that link for the overall G+C content of any genome (under NSB at equilibrium) as follows (**Supplementary Note S1**):

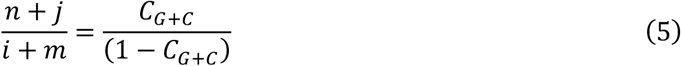

where the G+C content is naturally independent from *k*_*C*→*G*_ = *k*_*G*→*C*_ and *k*_*A*→*T*_ = *k*_*T*→*A*_ mutation rate constants. This results into a plot in **Figure 3c**, which shows where the G+C contents of different genomes should lie under NSB assumption at equilibrium.

The NSB assumption can lead to significant simplifications also when considering oligomers of size >1 for cross-mutations. For instance, the dyad case inter-mutational network can be represented by a tesseract (hypercube), with additional connections within, where each edge represents a mutation in one of the two bases between the dyad of one node to the dyad of another node (**Figure 3e**). We can see this on the example of the *ApC* → *ApT* mutation in **Figure 3d**. Under the NSB assumption, the system of 16 kinetic equations consisting of 96 independent mutation rate constants can be reduced into 48 mutation rate constants. The solutions show that this system, independently from the initial rate constant of base composition values, always equilibrates into a set of 10 unique solutions for the base content (**Figure 3f**, full derivation in **Supplementary Note S2**). Those 10 solutions, indeed, reflect PR-2, as follows:

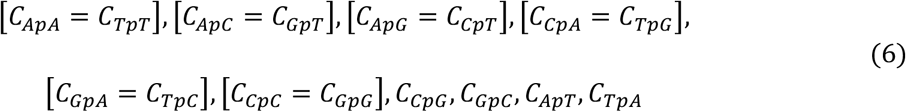

In an even simpler case, where we assume context independence of the mutation rates, the number of the independent rate constants can further go down into 3 unique solutions at equilibrium (analytical solutions found in **Supplementary Note S2**).

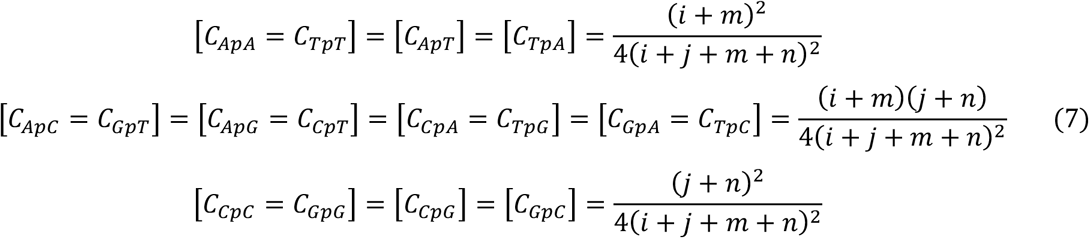

The above expressions reflect the connection between the genomic dyad content and the underlying individual rate constants. As expected, the three equilibrium solutions from dyad NSB, without context dependence of mutation rates, are the cross multiplications of the two unique equilibrium solutions obtained from singleton NSB, which is what is dictated by probability theory for the co-occurrence of independent events.

### 2.6. The plausibility of NSB assumption in actual genomes

Reviewed above, no strand bias (NSB) in the mutation rate constants (6,14) may become in conflict with experimental observations, as any deviation of the mutation rates from NSB, and/or genomic compositions from PR-2 would suggest that the model is wrong or that the system is out of equilibrium. In this section, we show that the NSB model can, overall, be considered as a good reductionist assumption for genomes, and highlight that there are indeed variations at a more regional scale and for some of our tested genomes. In **Section 2.5**, we showed that, under the NSB assumption, base compositional equalities at equilibrium is achieved and showed how this is linked with the overall G+C content of any genome. This results into a plot with an interesting behaviour shown in **Figure 3c**. To investigate how close the present day genomes lie along this ideal curve, we obtained the strand symmetric mutation rate constants from 17 species across the eukaryotic and prokaryotic kingdoms along with their G+C content and overlaid it with the curve in **Figure 4a** (30, 44–53). On average, the G+C content of the 10 eukaryotes deviate away by approximately 10% and 18% for the 7 prokaryotes. Interestingly, for all of the 17 species, except for *Aotus nancymaae*, the observed mutation rate constants suggest that the true G+C content should, in fact, be lower than what we observe in the present day. This may suggest that the genome composition is yet to reach the full equilibrium under NSB. Indeed, as we shall explain in further detail in **Section 2.8**, the NSB model suggests that the average time to reach PR-2 compliance occurs within 4.28 byr, the current maximum estimate for the age of life on Earth (43). However, using the strand symmetric mutation rate constants for 6 of the 10 eukaryotes to numerically calculate the equilibrium base content, we find that *Escherichia coli*, *Caenorhabditis elegans*, *Drosophila melanogaster*, and *Homo sapiens* should be closer to or already reaching PR-2 compliance under NSB, while *Aotus thaliana* and *Saccharomyces cerevisiae* are expected to achieve equilibrium much later (**Figure S7**). These results broadly match the deviation of the G+C content between the theoretical value at equilibrium and the true value. To investigate the plausibility of NSB assumption further, we employed the Trek methodology (29), in which we studied the 7-meric context-dependent germline mutation rates for the human genome. We found that the 12 sets of mutation rate constants for each of the plus and minus strands of the dsDNA, still averaged across the whole genomic span for each strand, are, in fact, highly correlated for the human genome (**Figure 4b**). This indicates that, while allowing for a regional deviations, NSB holds true for the average mutation rates summarised across the wider length of the human genome.

**Figure 4.**
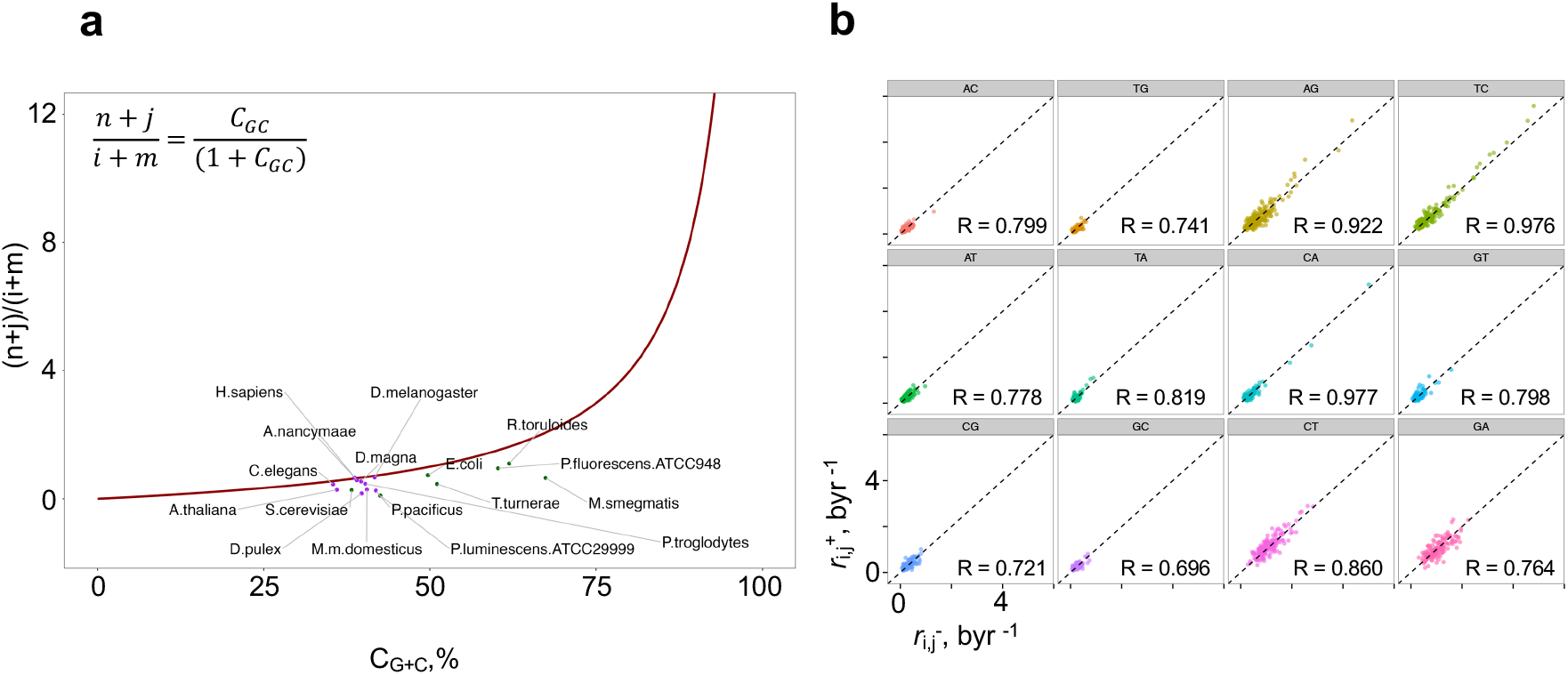
Strand-wise average mutation rate constants in humans. (a) The link between the mutation rates and the G+C content at equilibrium, along with the actual values for observed genomes (full details in **Figure S5**). (b) 12 sets of mutation rate constants, averaged across the length of the human genome, for each of the + and – strands of the dsDNA, obtained using the Trek methodology (22). The plots show the correlation between the 7-meric mutation rates in + and – strands, where each point represents one 7-mer with its central 4^th^ base undergoing the mutation of the plot category. The Pearson correlation coefficient values are indicated on each plot, where the kC➔A showed the highest value (R = 0.977).

### 2.7. A generalised link between mutation rates and PR-2

Since the prior considerations and our assumption-free simulation (**Section 2.4**) show substantial variation of the mutation rates that can result in PR-2 while not conforming NSB (**Figure 2**) Here, we tried to see whether a more generalised model can be found that can predict the PR-2 compliance of a genome, within 4.28 byr of evolution, not necessarily at compositional equilibrium, from mutation rates that are not necessarily NSB compliant. We used a symbolic regression modelling engine, Eureqa (now part of DataRobot), to find interrelations between the constituent 12 mutation rate constants from the sets under the tolerance region of PR-2 compliance (54,55). Eureqa, developed by Schmidt et al., determines the functional relationships to best describe the dataset in the simplest form *via* evolutionary search. The algorithm starts out with a random initial population of some unary (trigonometric, exponential, etc.) and binary (addition, subtraction, etc.) operators that collectively make up the tree-based graph. Next, evolutionary algorithm mechanisms take place, including mutation and crossover, each with an associated probability, to create offspring. A fitness score is evaluated (mean squared error) on these newly created offspring. The fittest offspring will be selected for reproduction, which, in symbolic regression, is evaluated by which functional forms describe the data better than others. The least-fit offspring will be replaced by a new population and this process repeats.

Each mutation rate constant was evaluated as a function of the remaining 11 mutation rate constants in the PR-2 compliance zone, which revealed more general mathematical dependencies that are more relaxed than the simple equalities of mutation rates under NSB assumption. Our new set of generalised mutation rate constraints can be described through sets of equations comprised of simple linear combinations of interchangeable mutation rate constants. For each mutation rate constant, Eureqa generated several equations, but we selected the ones that strikes a balance between low error (most accurate) and complexity (measured by the size and mathematical complexity of the symbolic expression). At times, the symbolic expressions could reach a high complexity, thus making it difficult to understand the interrelations between the mutation rate constants in a simple manner. We found that a complexity in the range of 7-9 terms provided this simplicity yet maintained its high accuracy. The found 12 simplified constraint equations are as follows:

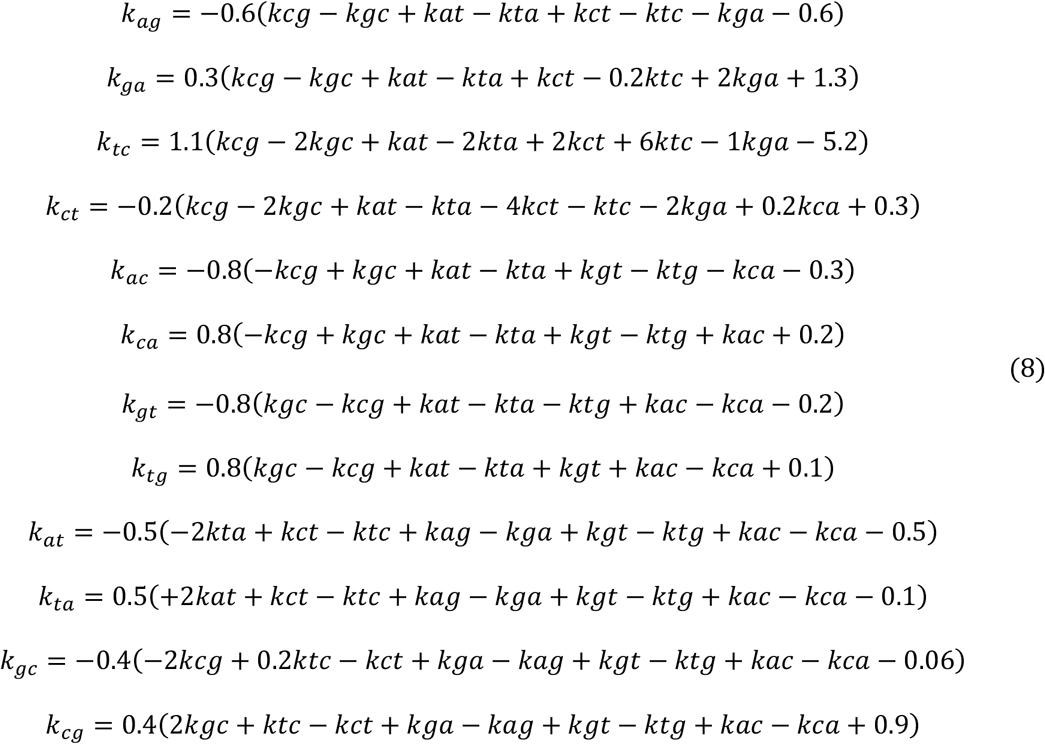

We can thus state that any mutation rate combination that complies with the above set of equations will lead to a PR-2 compliant genome. To test the performance of the 12 equations, we repeated the simulation of replicates following a non-symmetric, uniform distribution (equations in **Figure 1b**) by generating 10 million systems and applied the same PR-2 tolerance values from eukaryotic organisms. Using the test set on the 12 equations, we reveal that all equations show a strong, positive correlation between the true and predicted value (**Figure S6**). This shows that the equations are, on average, able to capture the majority of the mathematical dependencies amongst the mutation rates that result in PR-2 genomes with a high accuracy.

As initially discussed in **Section 2.6**, to examine how close the current PR-2 compliant lifeforms are to the fully equilibrated NSB solution (**Figure 5a**), we took the mutation rate constants of various species from the eukaryotic and prokaryotic kingdoms (30, 44–53) with their associated genomic G+C content and compared it to their expected NSB dependency at equilibrium (the theoretical line). We found that the majority of eukaryotic species are closer to the full equilibrium NSB solution, while prokaryotic species tend to be further away, despite all species falling within the PR-2 tolerance region (**Figure 5b**). Therefore, we can outline the presence of genomes that are far from NSB/equilibrium but are still PR-2 compliant. If we apply our set of 12 generalised constraints (**Eqs. 8**), we can verify that the mutation rates of such genomes are still within the relations to make a PR-2 compliant genome within 4.28 byr. Importantly, we believe this demonstration confirms that the drivers of PR-2 are still mutation rates, though with constraints more permissive and deviant from the previously proposed stricter NSB equalities. To convert the mutation frequencies into the genomic average mutation rate constants in a time domain (Trek-scaling), we used the equation *k* = 2.831*f* (see **Materials and Methods** for details on obtaining this equation), where *f* is the mutation frequency for a given mutation *k*_*s*→*u*_ for a given species. The resulting mutation rate constants were passed through the generalised equations above (**Eqs. 8**). Here, we demonstrate that the relations are linear and highly correlated (**Figure 5c**) for all the 12 constraint equations, more generalised than the simpler NSB equalities. To reiterate, **Eqs. 8** are obtained from a simulated system with no assumption and prior constraints on the mutation rates in our source simulations, but purely on the PR-2 compliance of the state after 4.28 byr of evolving. These equations are then applied to PR-2 compliant species with known mutation rate constants. Despite some of those species being out of the NSB-driven equilibrium, our relaxed equations still predict their PR-2 compliance. This reinstates the early view that behind PR-2 are the mutation rates, which we now show do not necessarily need to comply with NSB, as soon as they conform our more generalised constraints that seem to be in place for all the tried PR-2 compliant species.

**Figure 5.**
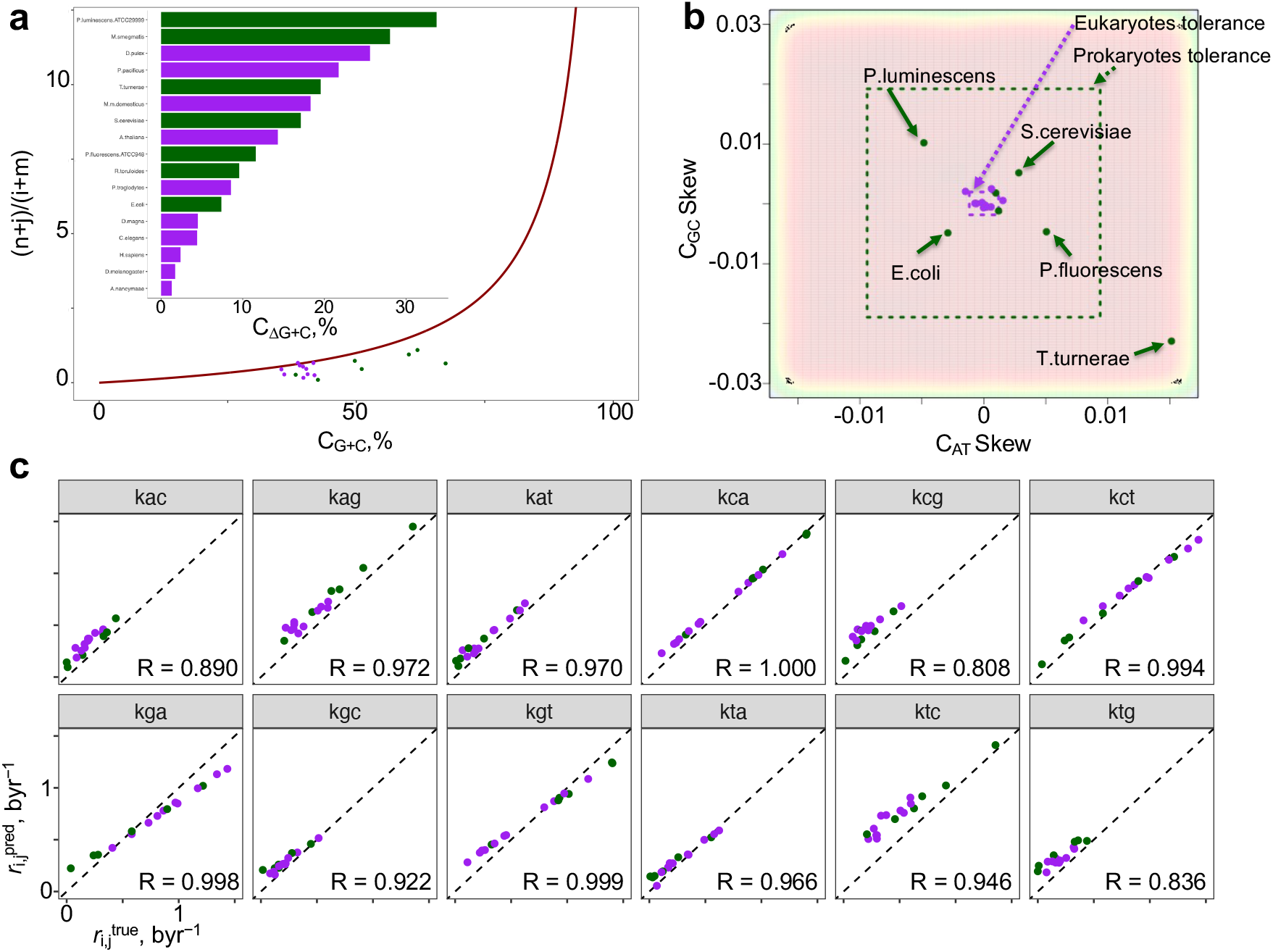
Comparison between species in terms of their closeness to NSB-driven equilibrium and closeness to the compliance to our generalised equations. (**a**) The base content solutions at equilibrium reveal the dependencies between the genomic G+C content and the mutation rate constants (red line). The strand symmetric mutation rate constants were obtained from 17 species across the eukaryotic and prokaryotic kingdoms (30, 44–53) and overlaid as a scatterplot with colourings based on their associated eukaryotic (purple) or prokaryotic (dark green) kingdoms. The difference in the G+C content between the true value and the theoretical value along the red curve was calculated for each species, coloured by kingdom, and represented as a bar plot within, sorted in decreasing order: *Photorhabdus luminescens ATCC29999, Mycobacterium smegmatis, Daphnia pulex, Pristionchus pacificus, Teredinibacter turnerae, Mus musculus domesticus, Saccharomyces cerevisiae, Arabidopsis thaliana, Pseudomonas fluorescens ATCC948, Rhodosporidium toruloides, Pan troglodytes, Escherichia coli, Daphnia magna, Caenorhabditis elegans, Homo sapiens, Drosophila melanogaster, Aotus nancymaae*. (**b**) The four systems of equations numerically solved for 25,000,000 systems based on the most general model of 12 independent mutation rate constants randomly drawn from a uniform distribution based on values obtained from the Trek methodology (29) in byr^−1^ range. Recording the base content GC skew and AT skew at the final 4.28-byr time point for all systems and zooming in to a AT skew range of ±0.015 and GC skew range of ±0.03, the 2-dimensional kernel density estimate scatterplot in **b** is obtained. The same 17 species from **a** are overlaid and coloured in the same scheme as in **a**. The dotted boxes represents the PR-2 compliant zones for the eukaryotic (purple) and prokaryotic (dark green) kingdoms, where each edge of the box represents the total 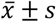 range for the corresponding axis (see main paper for exact values). (**c**) The strand symmetric mutation rate constants obtained from the 17 species across the two kingdoms, aligned to the Trek-scaling (29) and substituted the parameters of the 12 sets of mutation rate constant equations. Each plot represents one of the 12 mutation rate constants where each point represents a species, coloured by their corresponding kingdom in the same colouring scheme as in **a**. The Pearson correlation coefficient values are indicated on each graph, where the diagonal dotted line represents perfect correlation with a slope of 1.

The generalised equations (**Eqs. 8**) revealed above for each of the 12 mutation rate constants were evaluated as a function of the remaining 11 rate constants in the PR-2 compliance zone. Still focusing on the complete spectrum of mutation rate constant repertoire, we next explored a different approach. Can we have a machine learning model that can predict the PR-2 compliance from only the initial rate constants, without numerically solving the ODE system? To this end, we developed a machine learning model to classify PR-2 compliance as a function of the 12 independent, uniform distributed mutation rate constants (**Section 2.3**). The model revealed that the four mutation rate constants *k*_*A*→*T*_, *k*_*T*→*A*_, *k*_*G*→*C*_, and *k*_*C*→*G*_were the most important for classifying PR-2 compliance (see **Supplementary Note S2** for details).

### 2.8. General implications for Life: how long does it take to converge to PR-2?

Developing our simulations towards targeting more generalised characteristics of life, here we seek to explore the following three questions. 1) What is the average time it takes a genome to reach a PR-2 compliance? 2) What is the average time it takes a genome to reach a compositional equilibrium? 3) Are there significant differences between the times of (1) and (2). To do this, we analysed the kinetic system of equations by starting from random {*A*, *T*, *G*, *C*} contents within the range of allowed base content values in eukaryotic and prokaryotic organisms with symmetry-constrained mutation rate constants randomly drawn from a normal distribution. Previously, all simulations started with the initial base content of 25% for each of the four bases, however, in the following work, we remove this restriction as we seek to explain the universal nature of PR-2 from a species-invariant perspective. We used the PR-2 tolerance values obtained from the prokaryotic organisms because we learned from the simulation work that the number of systems that fall within the PR-2 compliance zone using the eukaryotic organisms is too small of a fraction from which we could draw any significant conclusions. We can, however, circumvent this problem by using the PR-2 tolerance values from the prokaryotic organisms instead and we also know that these species comply with PR-2.

Performing this simulation, we arrive to an average time to reach PR-2 compliance well within 4.28 byr (**Figure 6**), the current maximum estimate for the age of life on Earth (43). This happens independently from the initial genome composition, and from the randomly selected wide-range of mutation rates at around their human genome values. Interestingly, the system can start to comply with PR-2 even before reaching the base content equilibrium. In fact, on average, the PR-2 compliance reaches approximately one byr before the time to reach compositional equilibration (see **Materials and Methods** for PR-2 tolerance and genome equilibrium definitions). The average time to reach PR-2 compliance and genome equilibration becomes earlier as we increase the scaling factor. This may be explained in the following way: as we increase the scaling factor, we independently pick from a wider range of values from which one can randomly draw the mutation rate constants. As we do not consider the inter-dependence of the different mutation rate constants, we naturally arrive to a situation where the difference between the forward and reverse rate of reaction is greater, thereby arriving to an equilibration faster. These observations explain the species-invariant and universal nature of PR-2, as all the genomes have had enough time to reach the compliance.

**Figure 6.**
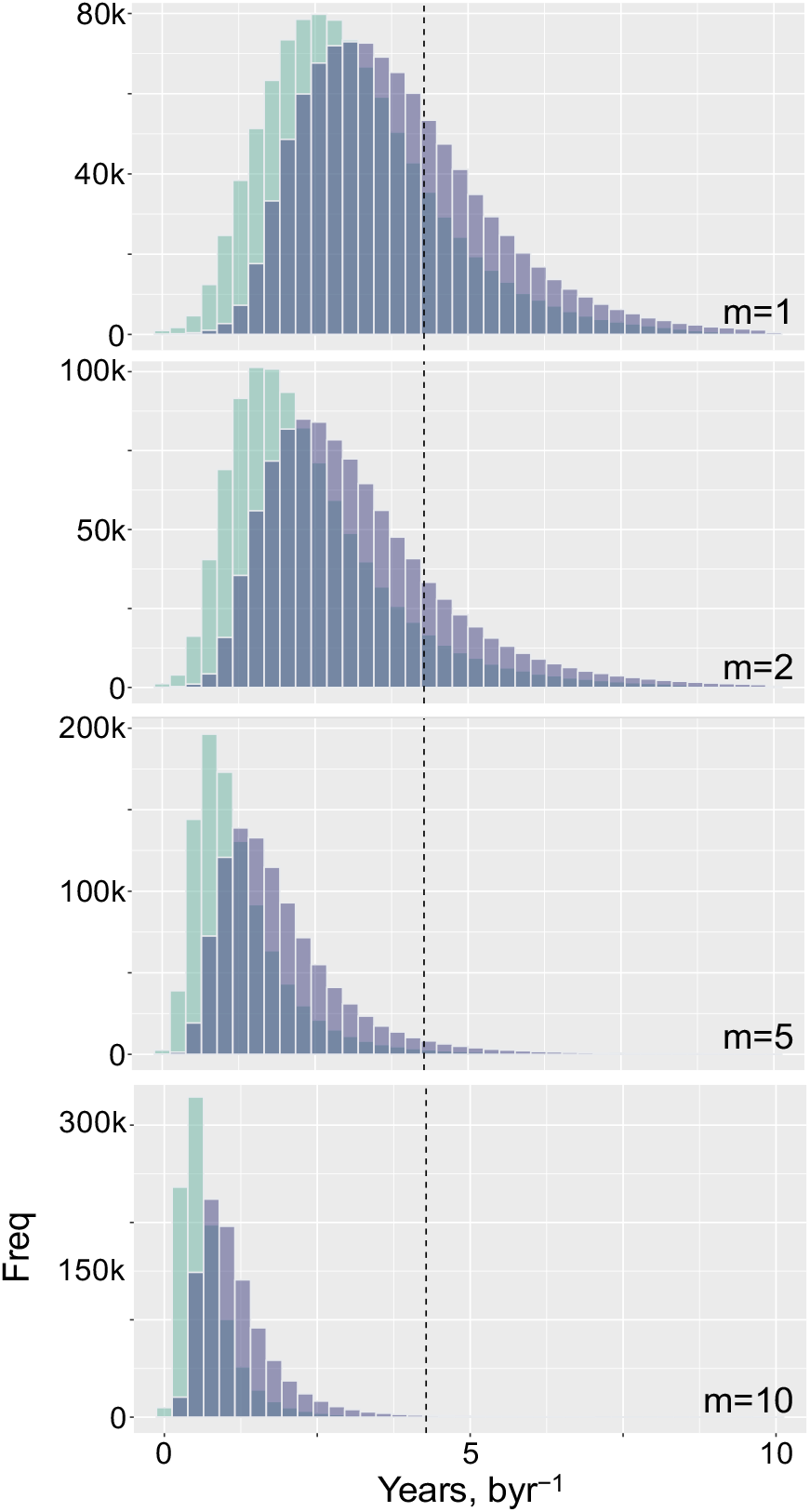
Distribution of time for a genome to reach PR-2 compliance and compositional equilibration. 10 million systems with the simulation model were generated where the initial base content was randomly sampled from the maximum allowed range based on prokaryotic organisms. The strand symmetric mutation rate constants were randomly drawn from a truncated normal distribution. This process was performed four separate times, each with a different multiplier, *m*, applied to the standard deviation of the random sampling of the mutation rate constants *m ϵ*{1,2,5,10}. The green histograms represent the distribution of time to reach PR-2 compliance and the purple histograms represent the distribution of time to reach genome equilibrium (see **Materials and Methods** for details). The vertical line intercepts the x-axis at 4.28 billion years, the maximum current estimate of age of life on Earth (43).

To compare this emergent behaviour to extant genomes, we obtained the strand-symmetry-accounted normalised mutation fractions from work done by Michael Lynch (30). These values first need to be converted to mutation rate constants in time domain (average mutation rate per site per billion years) so we can use it in our simulation model (see **Materials and Methods** for details). Following conversion, we set these mutation rate constants as the mean value in the normal distribution per species. As done previously, we generate 10 million systems with the simulation model for each of the six species. *Yeast* and *Arabidopsis thaliana* take, on average, the longest time to reach equilibration of the genome and PR-2 compliance. *Caenorhabditis elegans*, *Drosophila melanogaster* and *Escherichia coli* take a similar amount of time to reach equilibration and PR-2 compliance and all below the current maximum estimate of age of life on Earth (**Figure S7**). The difference between the time to reach PR-2 compliance and genome equilibration averages well below 2 byr.

## Abbreviations

The abbreviations used in the paper denote

dsDNA: double stranded DNA
ssDNA: single stranded DNA
NSB: no-strand-bias
PR-1: Chargaff’s first parity rule
PR-2: Chargaff’s second parity rule
myr: million years
byr: billion years
SNP: single nucleotide polymorphism
XGBoost: extreme gradient boosting
ML: machine learning
AUC: the area under the curve
ROC: the receiver operating characteristic
AUROC: the area under the receiver operating characteristic curve.

## DATA AVAILABILITY

The computer code, necessary to process the species across the three kingdoms, perform the simulations, employ the machine learning strategy perform the remaining computations, can be accessed through the following GitHub repository: https://github.com/SahakyanLab/GenomicPR2Simulations, and the ATGC Dynamics Solver web application is accessible on https://github.com/SahakyanLab/GenomicPR2SimsWebApp. All the used public datasets are accessible from the established genomic data repositories as detailed in **Materials and Methods**.

## ACKNOWLEDGEMENTS

PP is grateful to MRC, Hertford College, Clarendon Fund and Radcliffe Department of Medicine for supporting his DPhil studies. The Sahakyan Laboratory has been supported by the UK Medical Research Council (MRC), MRC Strategic Alliance Funding (MC_UU_12025).

## Supplementary Information

### Note S1. Derivations of the full equilibrium solutions

#### Note S1.1. The analytic solutions of the mutation rates under no-strand-bias assumption at equilibrium

Under the no-strand-bias (NSB) assumption, we can consider the example of k_C→T_ = k_G→A_, where the *s*→*u* subscript denotes the mutation of the base *s* into *u*. Here, k_s→u_ is the rate constant, which involves all processes that initiate and fixate the mutation in dsDNA, also converting the complementary strand. Central to our model is a plausible assumption of the strand-invariance of the mutation rates, i.e. the *s→u* substitution happens at the same rate independently from whether *s* is in the template or complementary strand of the dsDNA. To this end, at whichever rate C mutates to T in the template strand, with the same rate C converts to T in the complementary strand. Therefore, each strand will also have the complementary G to A conversions with the rate similar to C to T conversion, hence k_C→T_ = k_G→A_. This symmetry in rate constants significantly simplifies the mutation network from 12 to six independent rate constants, as such:

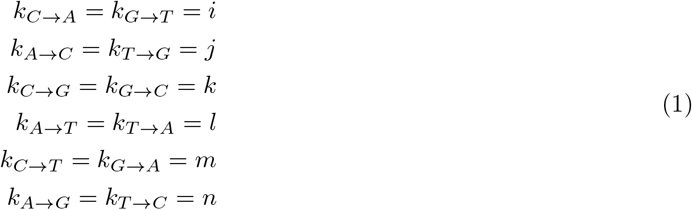

The reduced number of independent rate constants under the NSB assumption, produces a system of four kinetic equations, as such:

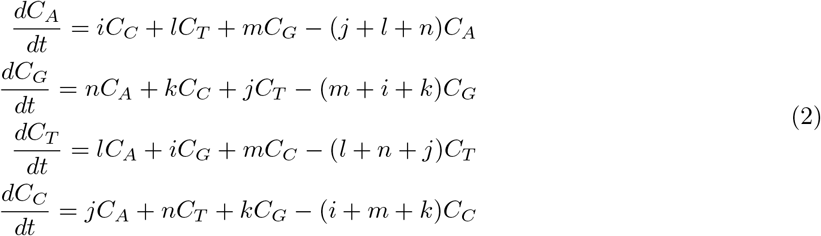

each describing the evolution of A, G, T and C base contents in fractions, making it possible to infer the time evolution dynamics of genomic base composition using different values for the rate constants and the initial base contents. Taking into account the outlined NSB-driven equalities, we can convert the system of four ODEs into a substitution matrix, *M*, as follows:

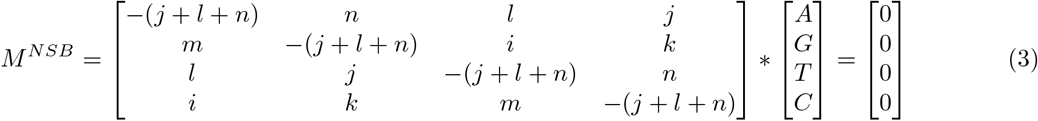

from which we can solve for the equilibrium base contents. In *Mathematica*, the system can be specified as:

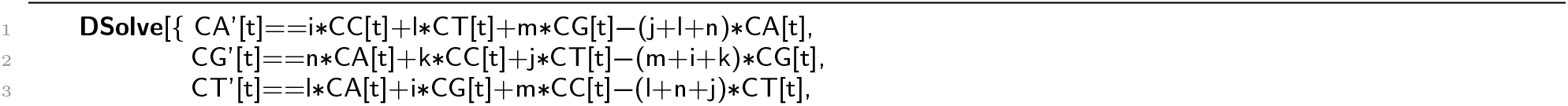

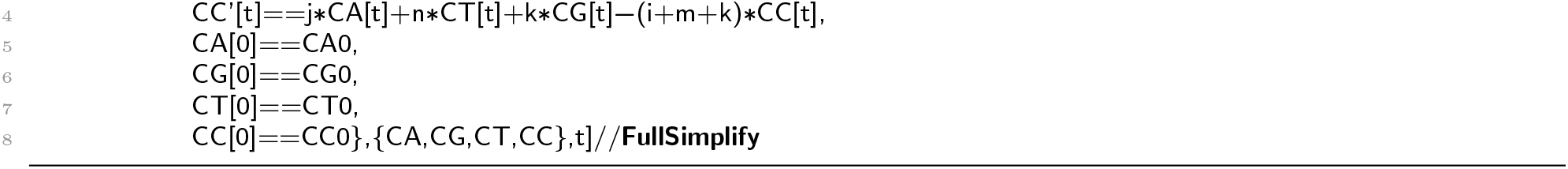

The solution to the ODE system is, however, too complex but we can obtain the symbolic solution of the system at equilibrium. There, the base contents are supposed to stay constants, hence we need to solve the system of equations displayed below, additionally setting the sum of all the base contents (in fractions) to 1.

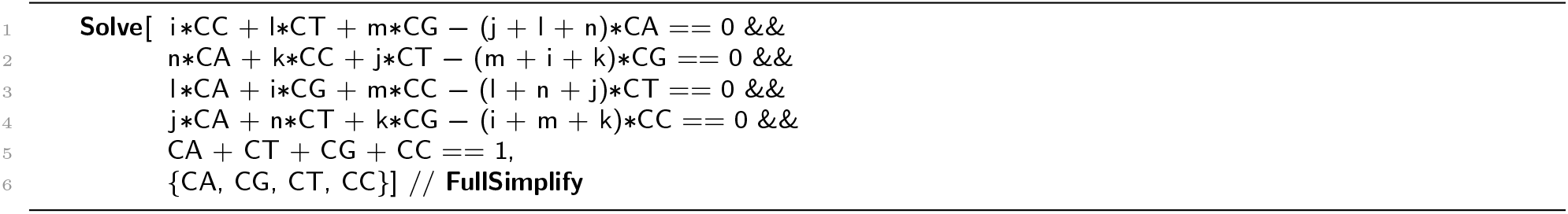

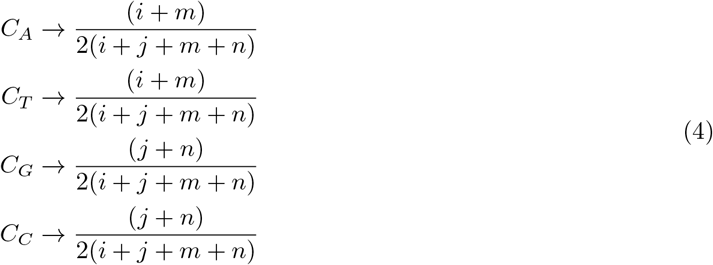

The system is fully solved, and the solutions imply the C_A_ = C_T_ and C_G_ = C_C_ equalities at equilibrium under NSB assumption for mutation rates. These solutions also link the mutation rate constants with the equilibrium genome composition. We can express that link for the overall G+C content of any genome, under the NSB equilibrium, as follows:

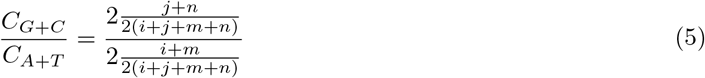

Since *C*_*A*+*T*_ = 1 − *C*_*G*+*C*_,

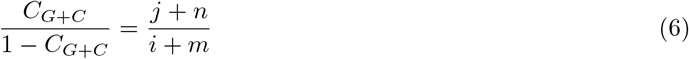

where the G+C content is naturally independent from k_C→G_ = k_G→C_ and k_A→T_ = k_T →A_ mutation rate constants.

#### Note S1.2. The NSB principle can be extended into higher k-meric orders

The NSB principle can lead to significant simplifications when considering oligomers of any size for cross-mutations, hence we can potentially exploit the extra information dyad-, triad-etc. counts may reflect on the genome. In section, the rate constants were reflecting the average singleton mutations across different sequence context found in a genome. Thus, the above model is applicable for finding the individual base contents in the scale of the entire genome. In contrast to this, when we want to recover the k-mer *k* ∈ {1,2,3,4,…} frequencies in a given genome, we need to consider the rate constants for many more k-mer transitions, as the neighbouring bases alter the mutation rate at a given site. To clarify this, consider the k_CpA→TpA_ and k_CpG→TpG_ substitutions in the dyad context. Both substitutions are a result of a C→T mutation. However, the rates for the two substitutions will be different (k_CA→TA_ ≠ k_CG→TG_), because of the neighbouring base (A vs. G) effects. In this particular example, we know that in the CpG context, the mutation rates for C are substantially elevated [1-4]. The equalities between the rate constants noted in section are true in the k-mer case as well, with the only difference being that we need to account for directionalities (5’-3’vs. 3’-5’) of the oligonucleotides. Hence, the rate constants of the substitutions that are reverse complementary to each other will be the same. We can see this on the example of ApT→ApT substitution.

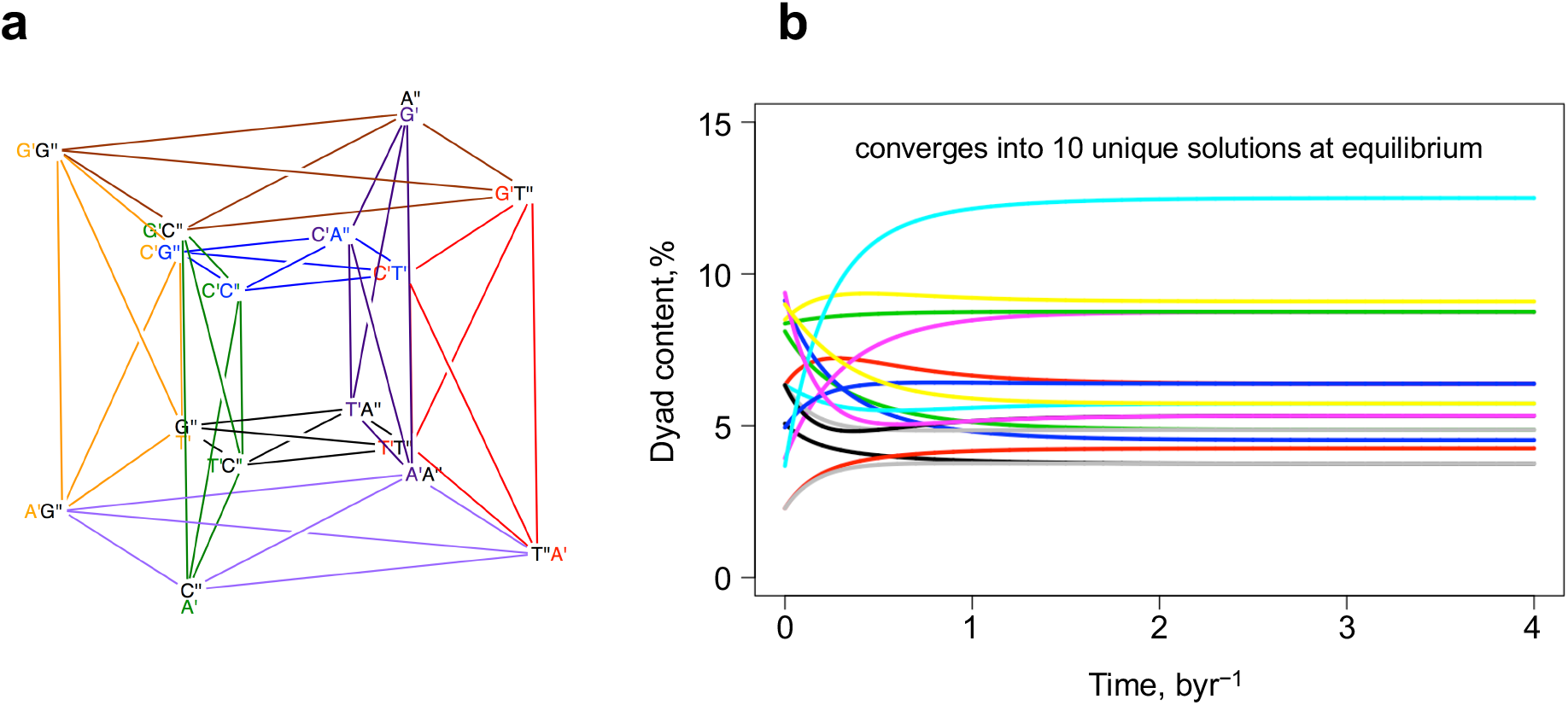
The NSB principle extended into dyads. The schematic representation of the cross-mutation network among dyads (**a**) is constructed based on the tesseract (hypercube). There, the primed superscripts denote the position in the dyad, from 5’ to 3’ direction (A’G” is the same as ApG). Each line should be interpreted as a set of two counter-directed arrows. The models, along with the reduction of the number of independent rate constants are shown below. **b** shows an example of the time evolution of dyad contents, modelled without and with context dependence for mutation rates, starting from arbitrary sets of initial dyad contents and rate constants. The colouring scheme is also random. The convergence to unique 10 dyad content values can be noted in **b**.

Taking into account that even the oligomeric mutations are prevalently driven by point mutations, [4] we can construct the network of cross-conversions between dyads on the basis of a tesseract (above Figure a), where the connections exist only between vertices that already have one common base. The superscript signs ’ and ” mark the first and the second bases, respectively, along the 5’-3’direction. Each line in the scheme should be interpreted as a set of two arrows in opposite directions. For this network, we can now construct the system of state equations by accounting for all the rate constant equalities, as described above for the dimeric case. The corresponding kinetic model is comprised of 16 equations for 16 unique dyads. In *Mathematica*, the system can be specified as:

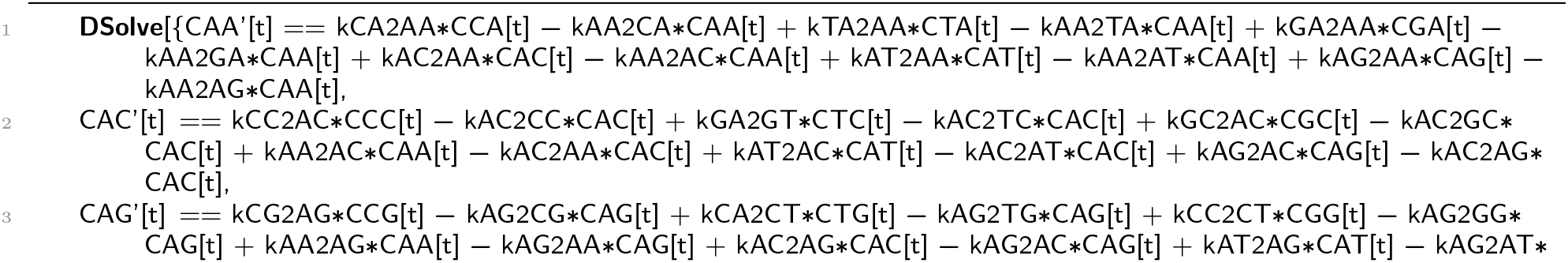

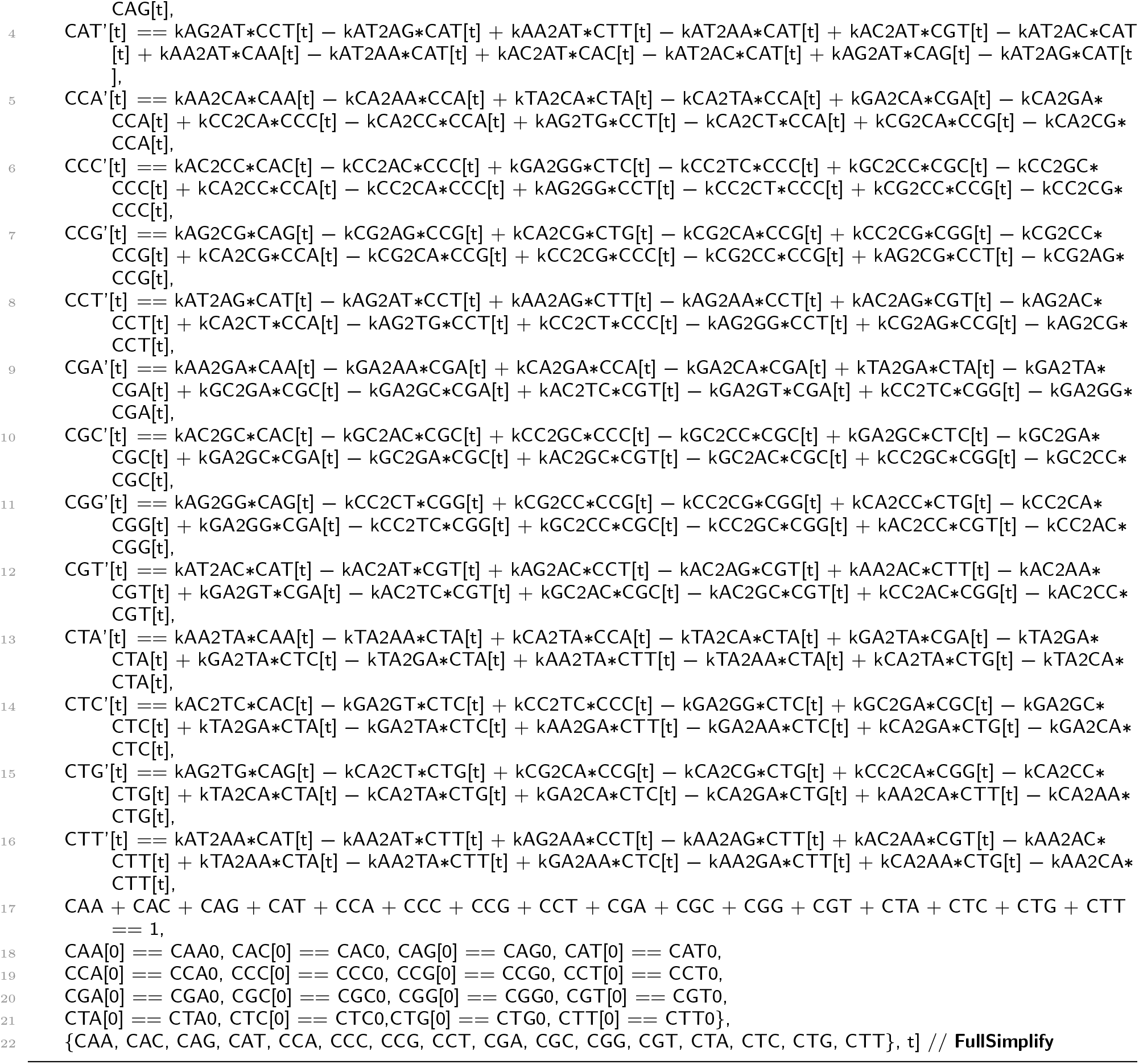

The equilibrium dimer fractions can therefore be computed *via*:

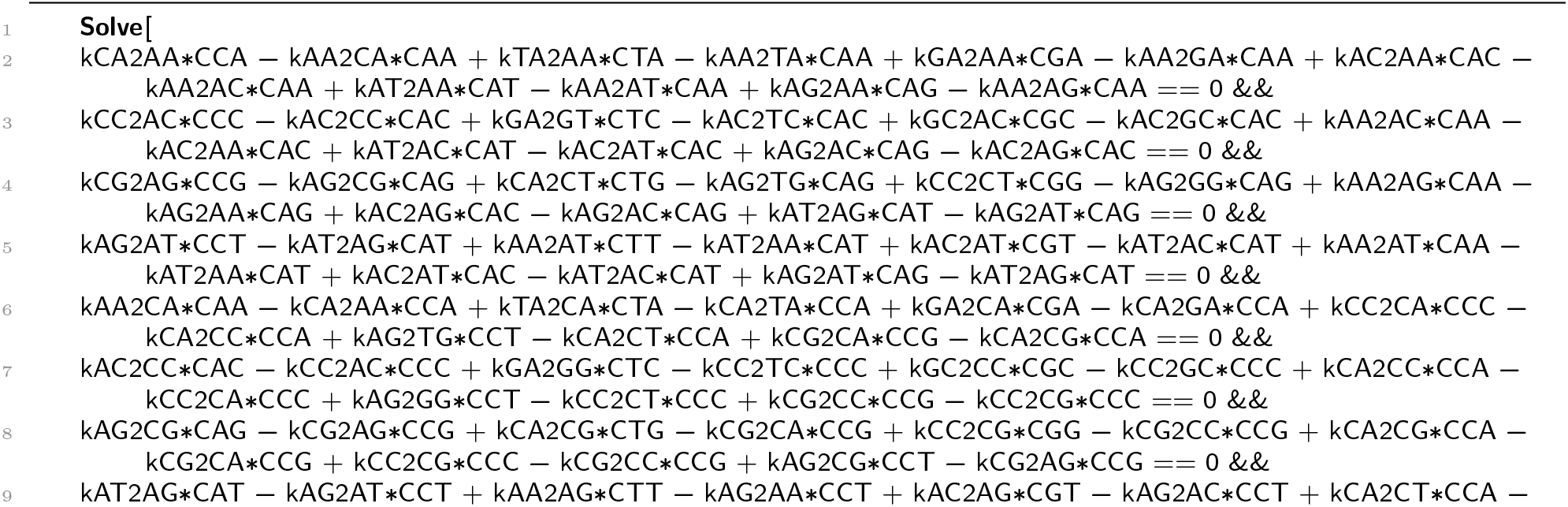

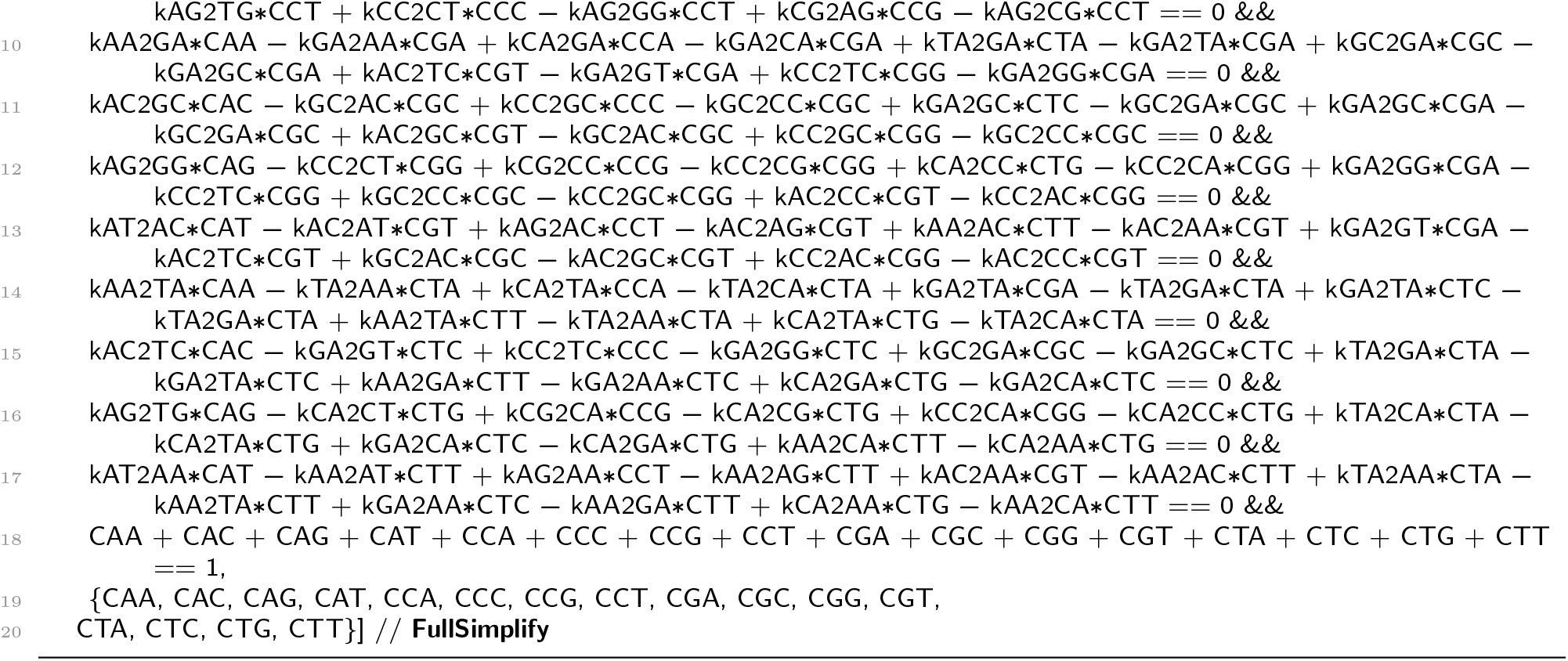

However, the above system of 17 equations with its 48 unique coefficients (unique after accounting for the rate constant equivalencies caused by double-strandedness of DNA) is very heavy for *Mathematica*. Adding different conditions, such as the rate constants being greater than 0, setting the dimer fractions being always greater than 0, restricting the solution region to different boundaries and trying all the matrix methodologies discussed above, does not help in getting solutions with 24GB of RAM and 12 CPUs. By exploring the solutions for an arbitrary set of numeric inputs, one can, however, see the equivalencies in the pairs of evaluated C_ij_ dimer contents that are reverse complementary to each other. This is exactly what the extended version of the Chargaff’s second parity rule states, which means that the oligo version of the observed species-invariant base-count equivalencies in genomes are also the result of the intrinsic rate constant constraints in the cross-mutation networks. Besides the numeric trials, the equivalency is clear from the comparison of the state equations for the reverse-complementary pairs. As an example, let us consider the state equations for the equilibrium fractions of ApA and its reverse complementary TpT dimers. After slight reorganisations, the equations for C_AA_ and C_TT_ correspondingly look like:

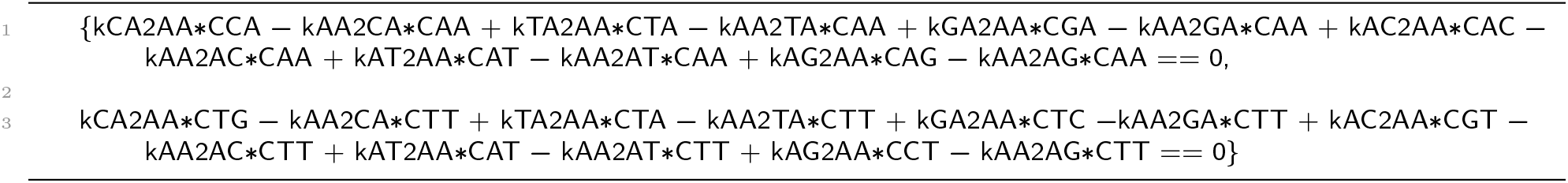

Those two equations have exactly the same set of rate constants and the same overall values of the expressions. Hence, one of the solutions to this pair would be the set where {C_CA_ = C_TG_, C_AA_ = C_TT_, C_GA_ = C_TC_, C_AC_ = C_GT_, C_AG_ = C_CT_}. Similar equalities can be inferred from the other 5 pairs of equations that have exactly the same set of rate constants (see below for the complete pairings).

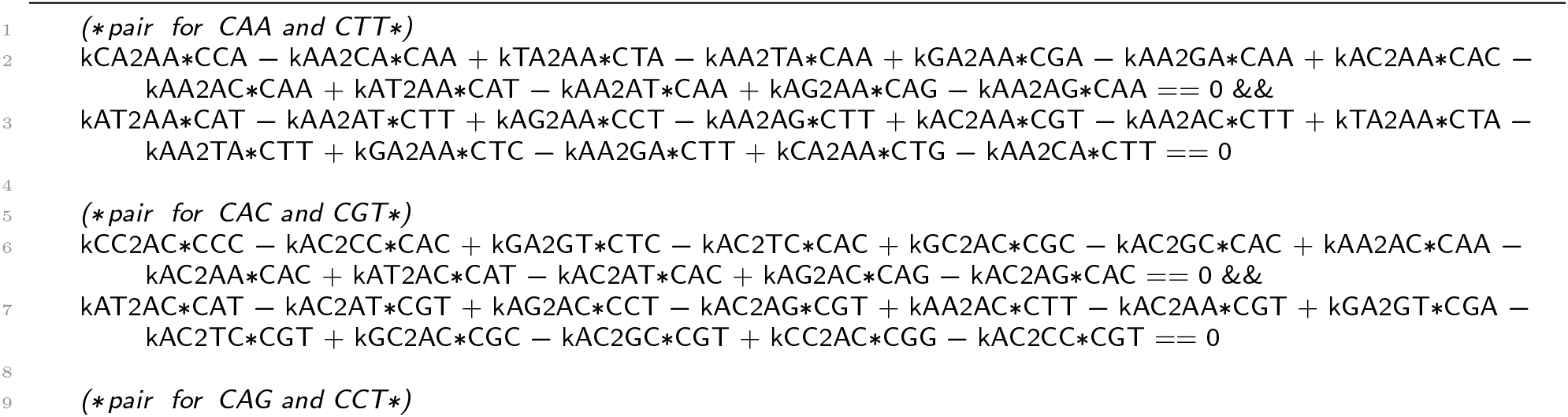

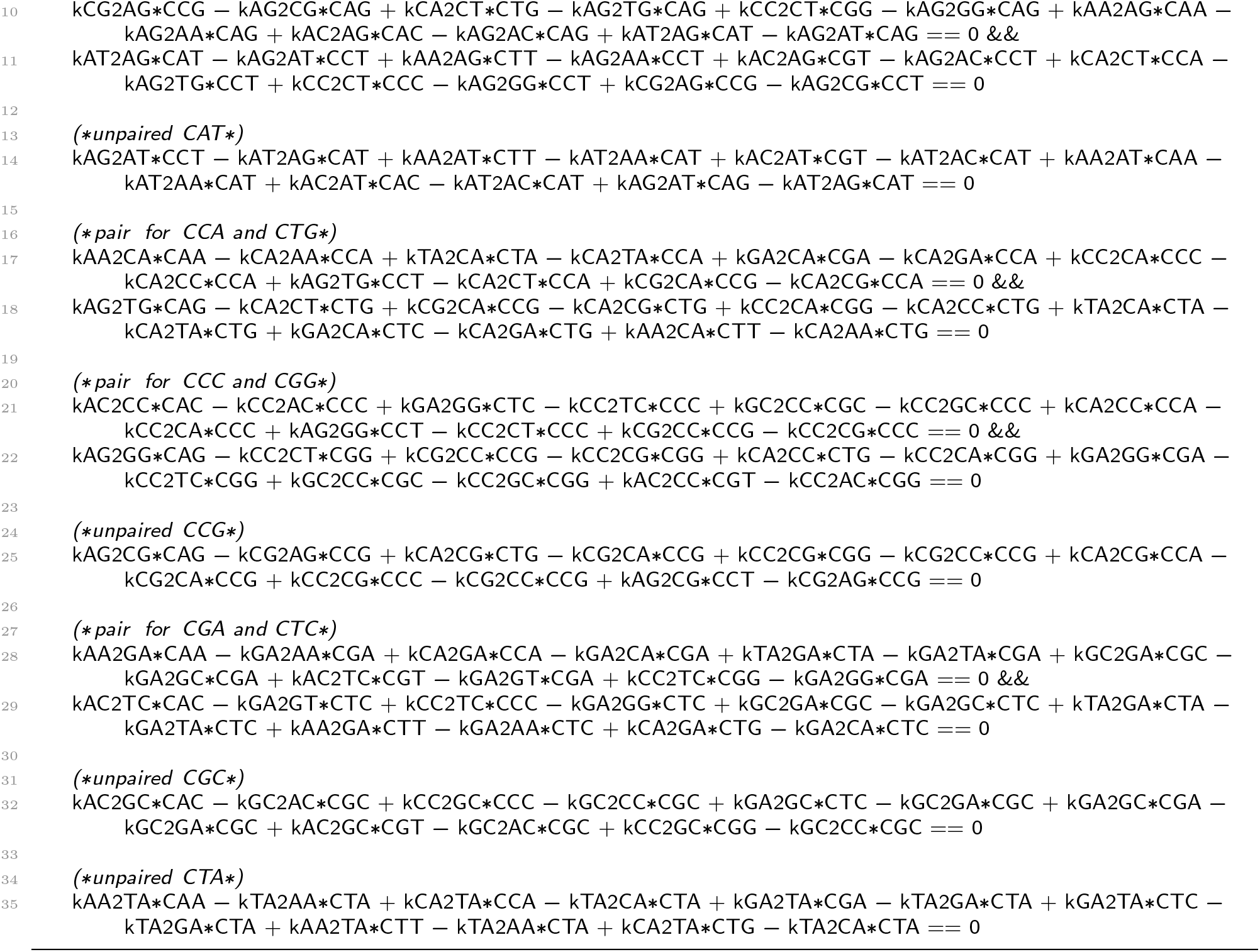

In summary, we have extended the cross-mutation network to represent the di-nucleotide (dyad) fractions in genomes. The set of rate constants in this extension are now accounting for the neighbouring nucleotide effects. However, we demonstrated that the rate constant symmetries, analogous to the single-base case, apply to the oligomeric extension as well. We then wrote down the system of differential equations *via* the reduced 48 mutation rate constants. Thus, the corresponding kinetic model, which is comprised of 16 equations for 16 unique dyads, always equilibrates into 10 unique solutions (above Figure b), fully complying with the oligo-version of PR-2, where the counts of the reverse complementary oligomers are also equal to each other in a single strand.

If we assume no context dependence for the mutation rate constants, we can use the same *i, j, k, l, m, n* rate constants from the 1mer-NSB model in section to describe all the cross mutations in the hypercube-based cross-mutation model. Of course, this assumption is crude, especially for some dyads where the neighbouring effect can be substantial (for instance the bases in CpG dyads have substantially higher substitution rates). However, this relatively reduced dimeric model may still be useful to describe the overall dyad contents in genomes and to reveal the magnitude of neighbour effects on substitution rates. The system of dyad-state equations therefore reduces into the following:

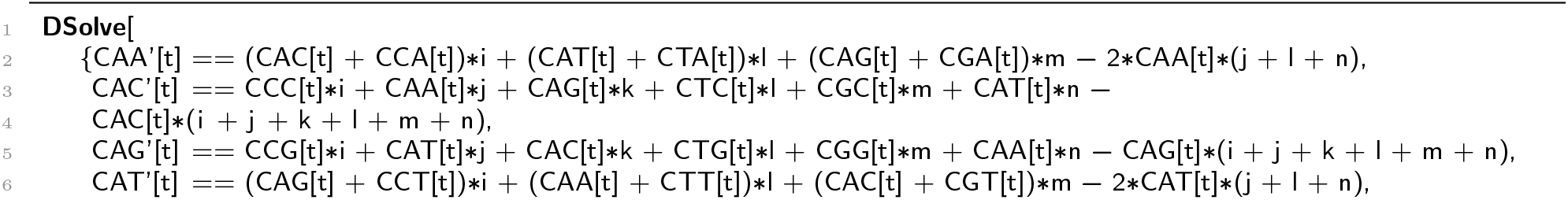

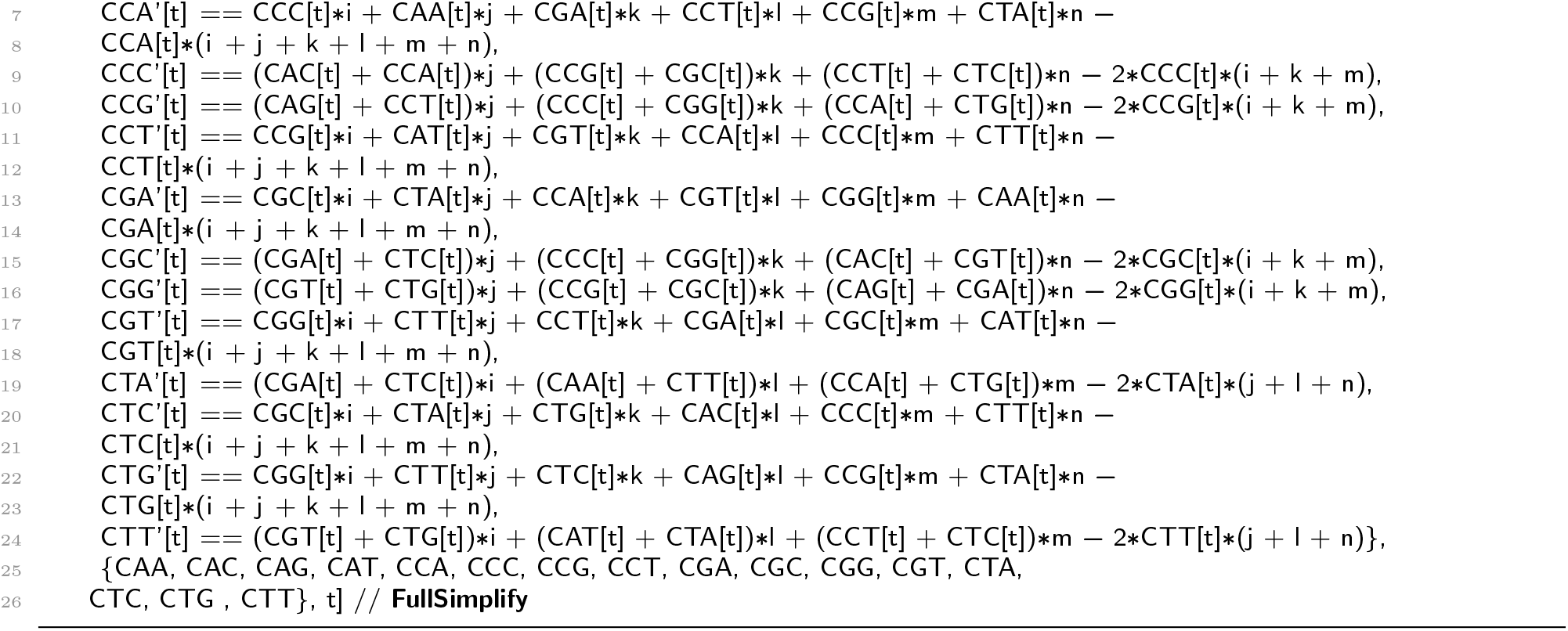

At equilibrium:

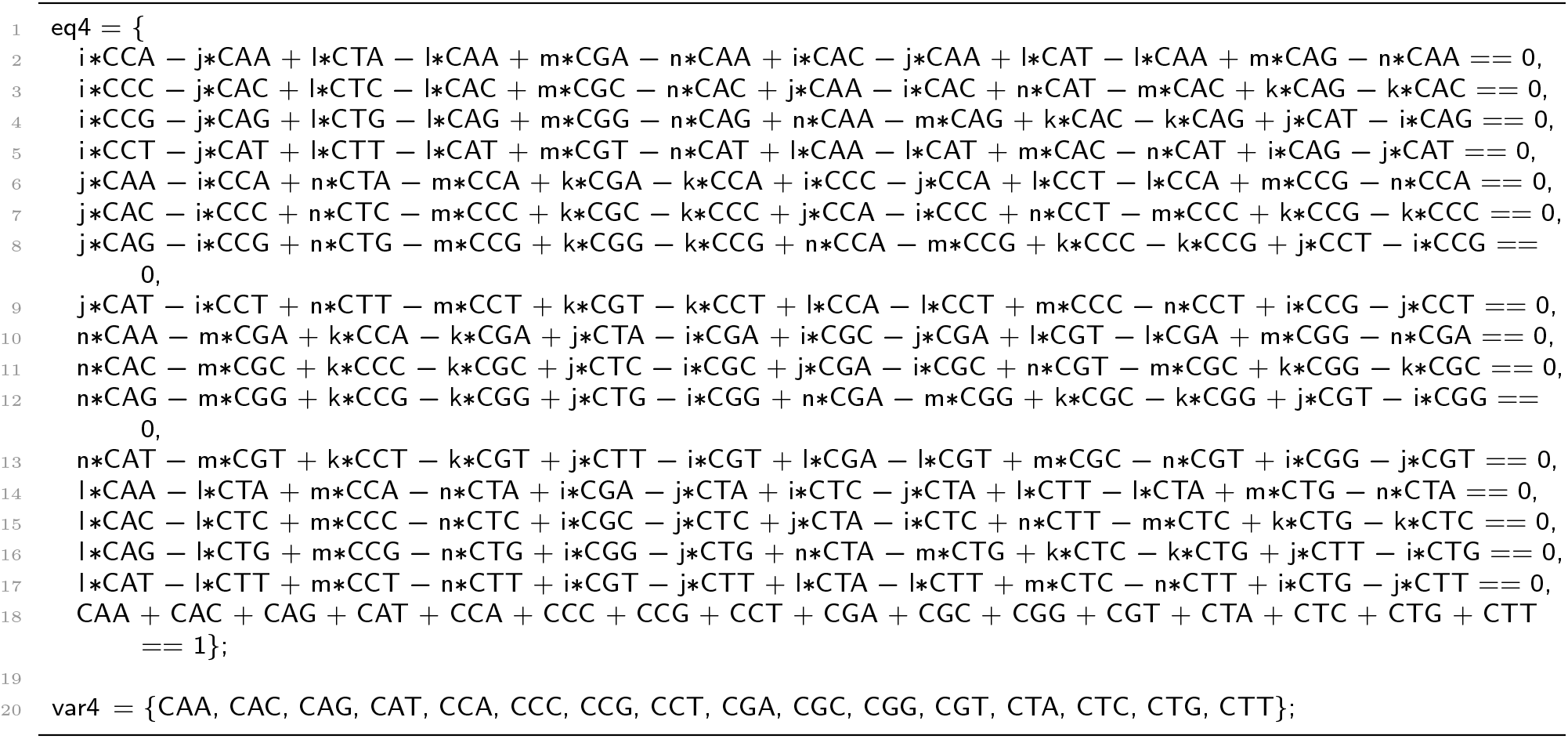

This system is still complex for *Mathematica*, but we can find exact symbolic solutions by carefully exploring the emergent groups in the equations. First, let us just assign an arbitrary set of numerical values to the {*i, j, k, l, m, n*} rate constants.

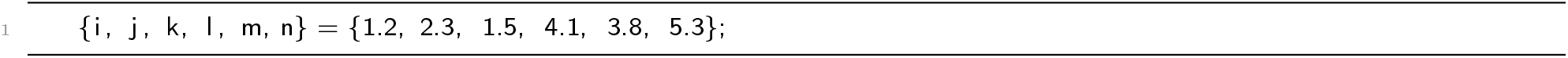

We can now find the numerical solutions:

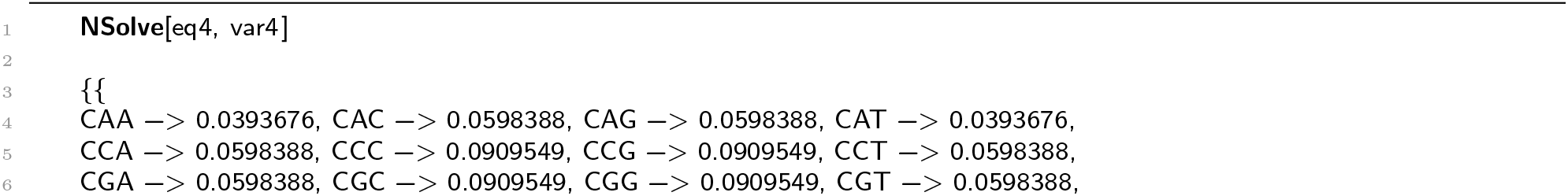

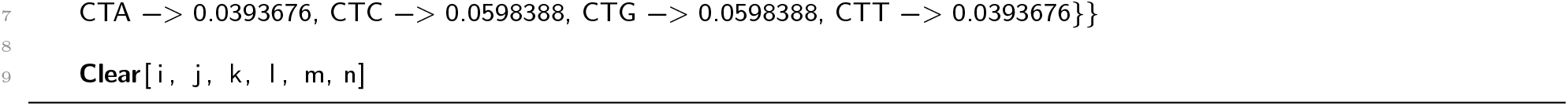

The numerical solution to the system with the arbitrary assigned coefficients implies that:

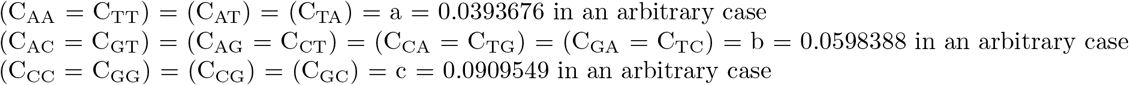

where, the brackets denote the equalities expected from the oligo version of the second parity rule. Hence, the empirical rule is, again, emergent from the solution of the system. We just have some additional equalities that are present under the assumption of no context dependence for the mutation rate constants. Therefore, for such hypothetical genomes, Chargaff could observe even more equalities, and the dimeric composition of the whole genome could have been described by just 3 values, *a, b* and *c*. Let us go on and use the found groups (invariant to the *i, j, k, l, m, n* numeric values, as soon as those are greater than 0) to further trim the system and get the symbolic solutions for *a, b* and *c*. The trimming of the system can be done by taking only one equation from each of the found three groups. The solutions to this model will be valuable as we can further compare the outcomes for real genomes with the actual dimeric contents, in which case the distortions from the theoretical values will reveal the presence of strong neighbouring effects on the mutation rates. We shall solve the reduced system by first creating the sparse arrays and solving the system of linear equations in a matrix convention. The solutions will be for the remaining C_AA_, C_AC_, C_CC_ dimer fractions that correspond to *a, b* and *c* above, representing the full solution to the complete system specified above.

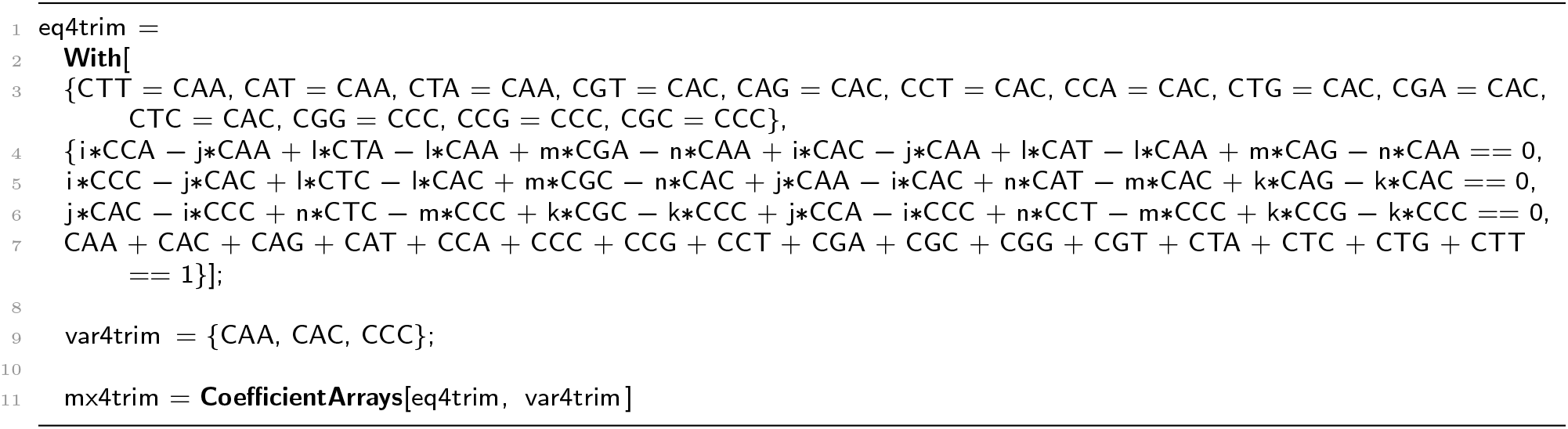

We can convert the above into a substitution matrix, *M*, as follows:

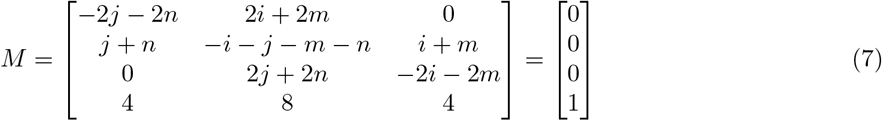

We can solve this equation in *Mathematica* with:

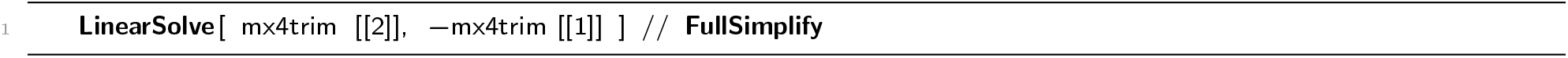

and obtain the following solutions:

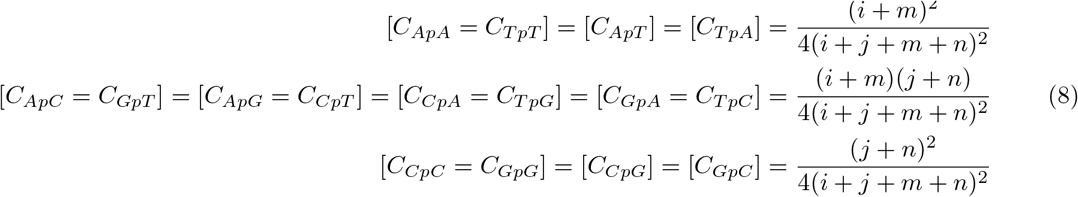

The unique {*a, b, c*} solution is found and can be easily verified by cross checking against the numerical solutions for any arbitrary set of non-0 and positive {*i, j, k, l, m, n*} rate constants.

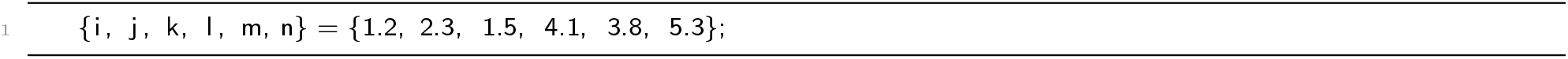

which returns the same values obtained numerically with **NSolve** above:

a = 0.0393676 in an arbitrary case
b = 0.0598388 in an arbitrary case
c = 0.0909549 in an arbitrary case

We have shown that, in this approximation, the model always converges into three unique dyad counts at equilibrium, the expressions of which reflect the connection between the genomic dyad content and the underlying individual rate constants. As expected, the three solutions from 2mer-NSB without context dependence for the mutation rate constants are the cross multiplications of the two unique solutions obtained from 1mer-NSB.

#### Note S1.3. Application of the mutation rate constants under no-strand-bias in predicating singleton and dyad composition of chimpanzee genome

Jiang et al. used genome-wide dSNP data from the chimpanzee genome to infer different substitution fractions [5]. The ancestral states of the sites were deduced by comparing the dSNP containing sequences to the homologous ones in humans. The work also reported the normalised substitution fractions, where each base is found in equal 25% frequency. Those substitutions are a result of the mutations happening within approximately 5-7 million years (*t*) after the divergence from the human-chimp most recent common ancestor [6, 7]. Since *t* is relatively small, we can assume that the probability of the sites undergoing repeated mutations is negligible. Since the data are normalised into a uniform base content, *n_i_* is always 0.25. Furthermore, both *n_i_* and *t* will cancel out in the context of the equilibrium base content equations derived above. To this end, the normalised substitution fractions can be used in a manner similar to the rate constants, as their scaled versions. Below we shall take the reported values, calculate the overall equilibrium base contents (both monomeric and dimeric) in the chimpanzee genome, and compare those with the actual genomic data. The SNP-sequence-context-normalised substitution fractions, in %, for the chimpanzee genome (from the described publication) are:

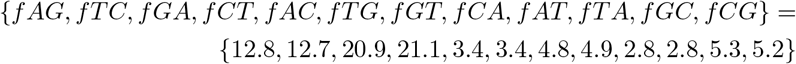

where *f_ij_* denotes the fraction of *i→j* base (single nucleotide) substitutions. We use these fractions as a replacement for the {*i, j, k, l, m, n*} rate constants. First, we average the already negligible differences between the reported substitution fractions, which are supposed to be equal by the NSB rate constant equality described in section.

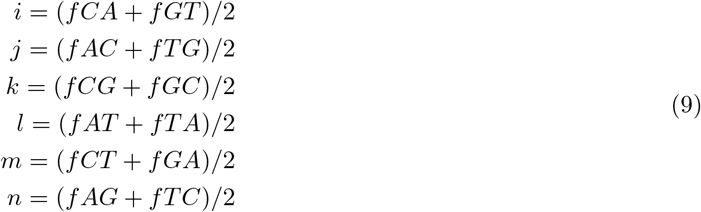

Now we can calculate the equilibrium base contents of the chimpanzee genome using the cross-mutation network solutions, where {*C_A_, C_G_, C_T_, C_C_* are the individual base contents, {*a, b, c*} are the dimeric contents under the assumption of neighbour-invariant rate constants, where

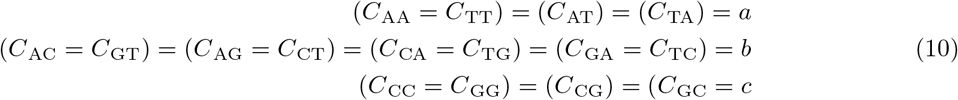

which results in the following individual base contents

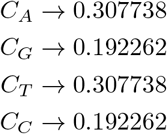

and the following dimeric contents

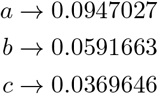

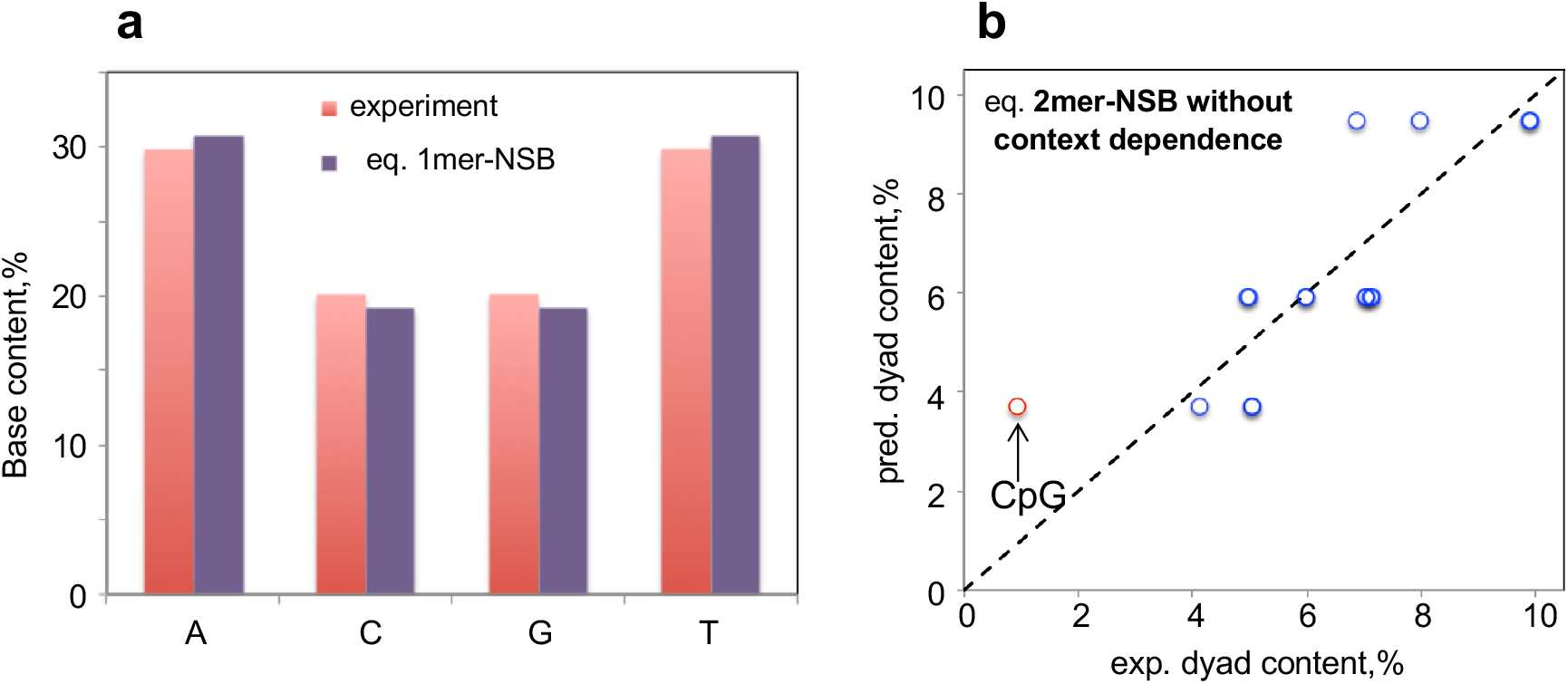
Application of the models on the chimpanzee genome. Comparison of the equilibrium single base contents, predicted via 1mer-NSB, with the experimental one extracted from the reference genome, **a**. Correlation between the dyad contents is shown in **b**, where the prediction is done via 2mer-NSB without context dependence for the mutation rate constants.

Unfortunately, we cannot apply the full hypercube model (without the neighbour invariance), which could have been applied numerically with NSolve in *Mathematica* if we would have the individual substitution fractions coming from 48 different dimer contexts. Comparing those with the experimental base and dyad contents of chimpanzee genome reveals a striking correlation. Since our solution is for the equilibrium genome, we can infer from the above Figure a, that the chimpanzee genome is slightly off equilibrium and will move towards it by having more G:C|C:G→A:T|T:A mutations, consistent with the prior view on the state of the chimpanzee genome [5]. As for comparing the predicted dyad contents, since we do not have the substitution fractions for the 48 independent dyad crossings to use those for the complete dyad inference (2mer-NSB), we have only used the solution to the hypercube model with the assumption of 2mer-NSB without context dependence for the mutation rate constants by using the six rate constants inferred from the single-nucleotide-based substitution data. The agreement is still rather impressive (above Figure b), where, as expected, the neighbour effect distorts the CpG dyad frequency the most since the bases in CpG have much higher mutation rates owing to the involvement of epigenetic mechanisms. We expect these differences to vanish, as better-quality substitution data become available, accounting for the sequence context and enabling the usage of the full 2mer-NSB model for dyads.

#### Note S2. Machine learning model for classifying PR-2 compliance

Here, we developed a machine learning model to the non-symmetric, uniform distribution simulation outcome to fit a classifier for compliance and non-compliance with the PR-2 solutions. This would give an impression on the full potential of the mutation rate constants to proxy mirror the PR-2 compliance status without actually solving the ordinary differential equation systems. To do this, we used the tree-based extreme gradient boosting (XGBoost) machine learning model as our central framework for the model development [8, 9]. In gradient boosting, an ensemble of learners is developed with each iterative learner predicting the residual of the ensemble of prior learners. The combination of the decision trees (maximum interaction depth and minimum child weight) as the underlying learner with the gradient boosting process (number of boosted trees, learning rate, subsample percentage and gamma) allows flexible tunability and optimisation of the six hyperparameters; a combination that is commonly used in a wide range of machine learning competitions such as Kaggle (www.kaggle.com) and predictive modelling [8, 10]. Another important reason for selecting this machine learning strategy is that XGBoost allows the extraction of information regarding the importance of features, thereby giving us insight into which of the 12 mutation rate constants are most important to predict PR-2 compliance.

We employed the XGBoost machine learning model as the central framework for the model development in our study available in R through the caret package [11]. For the machine learning model evaluation, we used the receiver operating characteristic (ROC) performance metric (using the MLeval package [12]) because, as we will describe in the following paragraph, our observations in the training set are balanced between each class. If it were skewed, other performance metrics may be more appropriate, such as the precision-recall curve [13–15]. We applied a k-fold cross validation (KCV) for evaluating the model (see Materials and Methods for details on hyperparameters). In a KCV procedure, the data set is randomly split into k smaller parts in order to reduce the risk of any over-fitting, and is a common practice for training machine learning models in the scientific literature [10, 16–19]. For instance, if k=5, the training data is split into 5 smaller sets, where, in each iteration, the hyperparameters of the model are trained using k-1 of the folds as the training data set. Next, the model is validated on the remaining part of the data set. Specifically, the remaining fold is used by the trained model as a test set to evaluate the performance metric, which, in our model, is the accuracy. This process is repeated k times, where, at the end of this process, the performance measure is reported as the average of the values in each k-fold. Given the size of the training data and common range of values for the k in KCV in the scientific literature, our model was trained with k=6 repeated once.

Using the tolerance values from eukaryotic organisms on the 25 million systems generated from the non-symmetric, uniform distribution simulation, we naturally end up with a disproportionate amount of non-compliant cases over the compliant cases. For the purpose of the machine learning process, we, therefore, used an equal number of compliant and non-compliant cases for the training set, starting from all 12 mutation rate constants as features. We also performed a principal component analysis on these 12 mutation rate constants for the equilibrated cases without imposing the PR-2 tolerance and found that there was no outstanding principal component (data not shown). The selection of non-compliant cases is done via random sampling (*seed* = 2022). We normalised the selected data set (*seed* = 1234) to randomly sample the seeds for each of the six cross-validation processes, repeated once. In the model training process, we tuned the learning parameters using a grid search approach. The performance metric selected for the development of the machine learning model was the receiver operator characteristic (ROC) curve, which shows the True Positive Rate (TPR) vs.

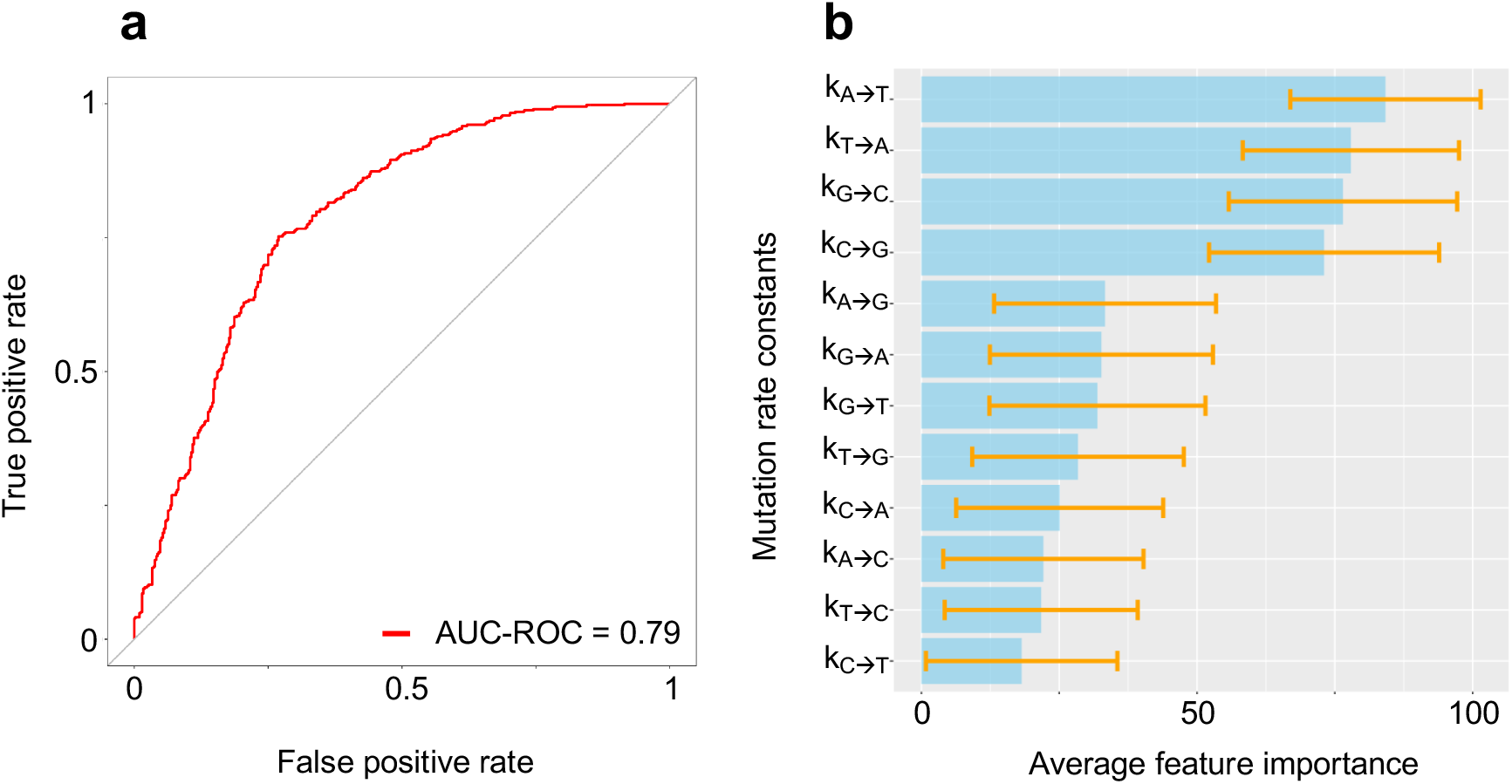
The performance of the final machine learning model strategy. (**a**) Area under the receiver operating characteristic (AUC-ROC) curve showing the performance of the classifier for compliance and non-compliance with the PR-2 solutions. (**b**) The 12 mutation rate constant-based features ranked by relative average importance for the achieved prediction quality. The selection of non-compliant PR-2 cases was done via random sampling. As the model is trained on a random selection of non-compliant PR-2 cases, the feature importance of the 12 mutation rate constants would not be representative of the wider data set. Instead, we repeated the training process 1000 times using the optimised hyperparameters to obtain an average feature importance plot of the 12 mutation rate constants. The barplots represent the average feature importance and the orange bar represents the average ± 1 standard deviation.

False Positive Rate (FPR) at varying classification thresholds. TPR is defined as the following:

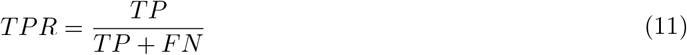

and FPR is defined as the following:

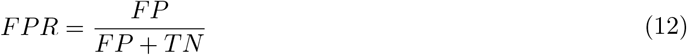

To quantify the performance of the model across all possible classification thresholds, we compute the Area Under the ROC Curve (AUROC). The feature importance was calculated using the varImp function from the caret package. The optimised model obtained an AUROC value of 0.79 (above Figure a). Given the model is trained on a random selection of non-compliant PR-2 cases, the feature importance plot of the 12 mutation rate constants would not be representative of the wider data set. To circumvent this problem, we repeated the training process 1000 times using the optimised hyperparameters to obtain an average feature importance plot of the 12 mutation rate constants (below Table). For each random sampling of non-compliant PR-2 cases, we fixed the seed to 123. The diagonal transversion mutation rates (k_A→T_; k_T→A_; k_G→C_; k_C→G_) have a consistently higher feature importance compared to all other mutation rates (above Figure b).

Key hyperparameters tuned for the eXtreme Gradient Boosting (XGBoost) classification model.

| **Hyperparameters** | **Description** | **Tried values** | **Optimised model** |
| --- | --- | --- | --- |
| Nrounds | Number of boosted trees to fit. | 200, 500, 1000, 2000, 3000, 4000, 5000, 7000, 9000, 11000, 13000, 15000, 17000, 20000 | 17000 |
| Max depth | Maximum depth of each tree. | 5, 6, 8, 10 | 6 |
| Gamma | Controls when to split a node. | 0, 0.1, 0.2 | 0 |
| Min child weight | Minimum sum of instance weight needed in a child. | 1, 5, 10 | 1 |
| ETA | Controls the contribution of each tree on the outcome and how quickly the algorithm learns. | 0.005, 0.01, 0.02, 0.1 | 0.01 |
| Subsample | Fraction of observations to be randomly selected for each tree. | 0.3, 0.4, 0.6, 0.8 | 0.3 |
| Colsample bytree | Subsample ratio of columns when constructing each tree. | 1 | 1 |

## Supplementary Figures

**Figure S1.**
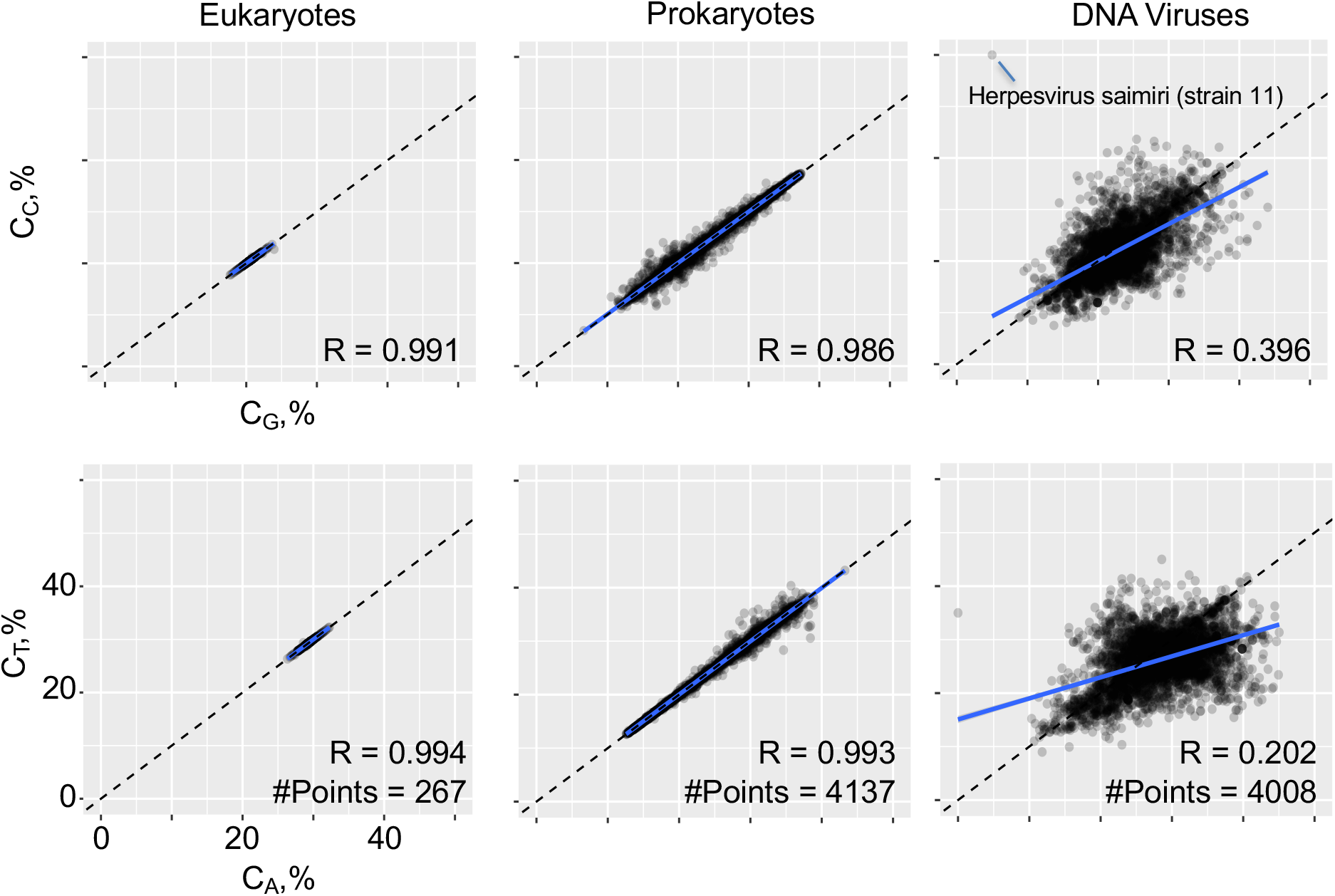
The percentage of guanine (G) vs. cytosine (C) and adenine (A) vs. thymine (T) computed for species in the eukaryotes, prokaryotes and DNA virus kingdoms. Each of the base contents are computed on the reference strand for a given dsDNA genome. (**top row**) Percentage of the G vs. C contents represented as scatterplots. Eukaryotes have the highest Pearson correlation coefficient value (R = 0.991), prokaryotes similarly high (R = 0.986) while DNA viruses exhibit the lowest correlation coefficient value (R = 0.396). Following the filtering process for the species in each kingdom (see **Materials and Methods**), the genome of the herpesvirus saimiri (strain 11) has the most extreme base contents compared to the other species in the DNA virus kingdom. (**bottom row**) Similar graphs were generated for the percentage of the A vs. T contents. Eukaryotes have the highest Pearson correlation coefficient value (R = 0.994), prokaryotes similarly high (R = 0.993) while DNA viruses exhibit the lowest correlation coefficient value (R = 0.202).

**Figure S2.**
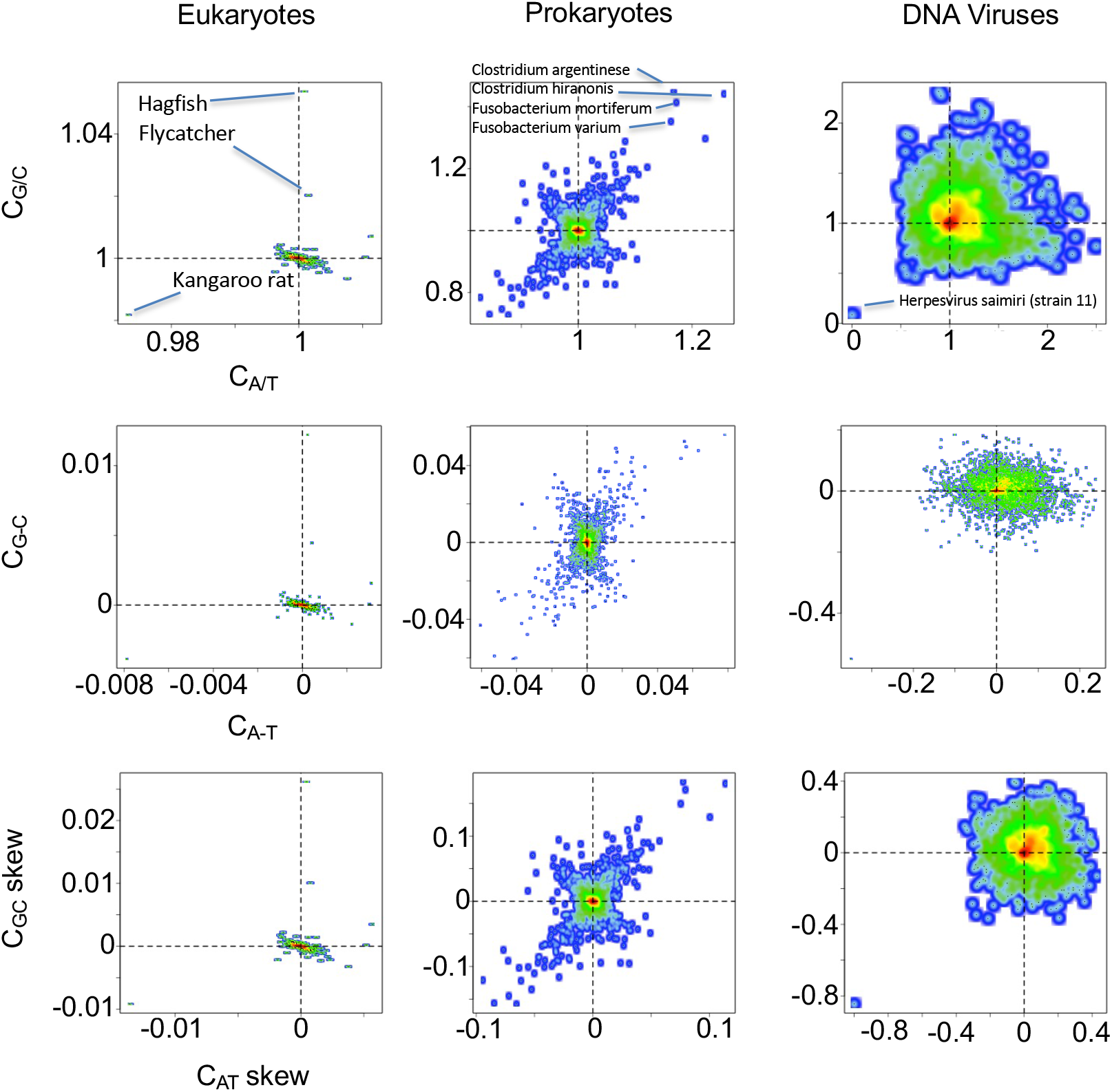
Nucleotide compositions for species in the eukaryotes, prokaryotes and DNA virus kingdoms. Each of the base contents are computed on the reference strand for a given dsDNA genome and represented as a 2-dimensional kernel density estimate scatterplot where the dotted vertical and horizontal lines indicate perfect parity. (**top row**) The base content ratio of C_G/C_ vs. C_A/T_ of eukaryotes have the tightest range with 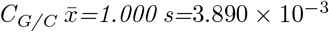 and 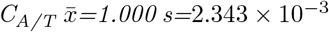. Prokaryotes are more dispersed compared to eukaryotes with 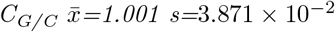 and 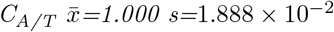. DNA viruses are the most dispersed with 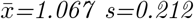 and 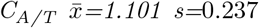. Following the filtering process for the species in each kingdom (see **Materials and Methods**), we highlight some species that have the most extreme nucleotide compositions for each of the three kingdoms. (**middle row**) The base content difference of C_G-C_ vs. C_A-T_ of eukaryotes are the smallest with 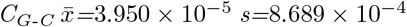 and 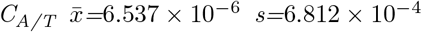. Prokaryotes are more dispersed with 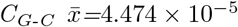 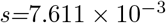 and 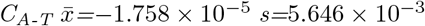. DNA Viruses are the most dispersed with 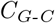 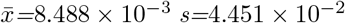 and 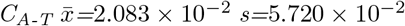 (**bottom row**) The base content C_GC_ skew vs. C_AT_ skew of eukaryotes have the smallest C_GC_ skew 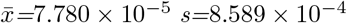 and C_AT_ skew 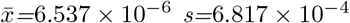. Prokaryotes have a C_GC_ skew 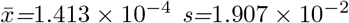 and C_AT_ skew 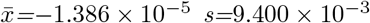. If the data set of the prokaryotic organisms is split by the average G+C content of 51.16% and plot the C_GC_ vs. C_AT_ skew separately for the G+C content above and below the average, we observe the present diagonal pattern (bottom left to top right) for below average G+C content, while a diagonal pattern (top left to bottom right) for above average G+C content. DNA viruses have a C_GC_ skew 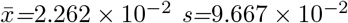 and C_AT_ skew 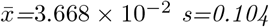.

**Figure S3.**
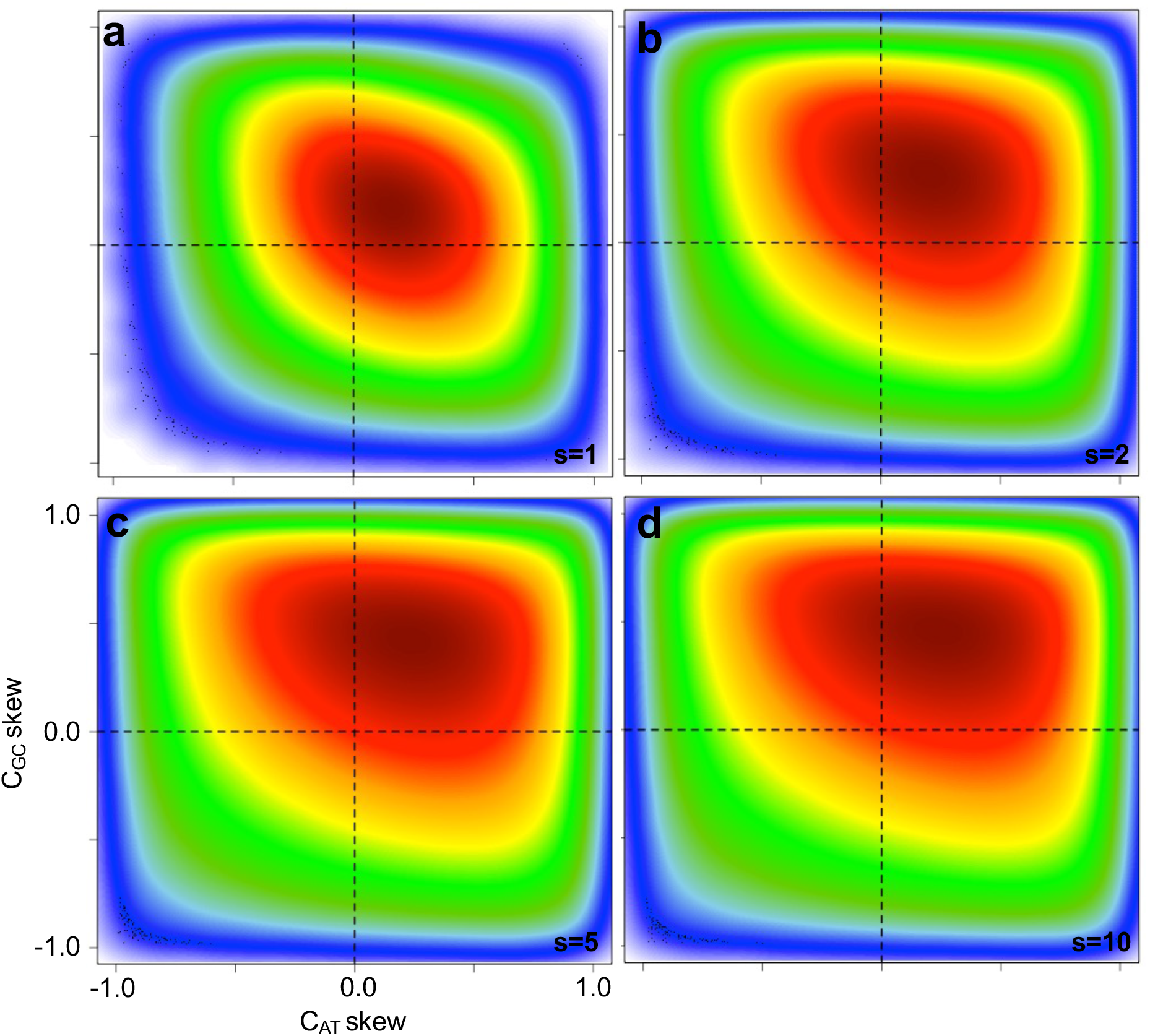
Numerical analysis of the NSB model with all-independent mutation rate constants assumed to have a normal distribution. The systems of equations were solved starting from 25% initial contents for all four bases and rate constants randomly and independently drawn from a normal distribution obtained from the Trek methodology [16] in byr-1 range. 25,000,000 such systems were calculated to produce genomic base contents at the final 4.28-byr time point. The 2-dimensional kernel density estimate scatterplot in c presents the distribution of C_GC_ and C_AT_ content skews from the outcome of the simulation (colours vary with decreasing occurrence frequency from red to blue), with the white dotted box indicating the 1.5x zone of compliance for the DNA virus kingdom with PR-2. The simulations were run four times separately with standard deviation values scaled by an additional multiplier m ∈ {1,2,5,10} in plots a-d, respectively.

**Figure S4.**
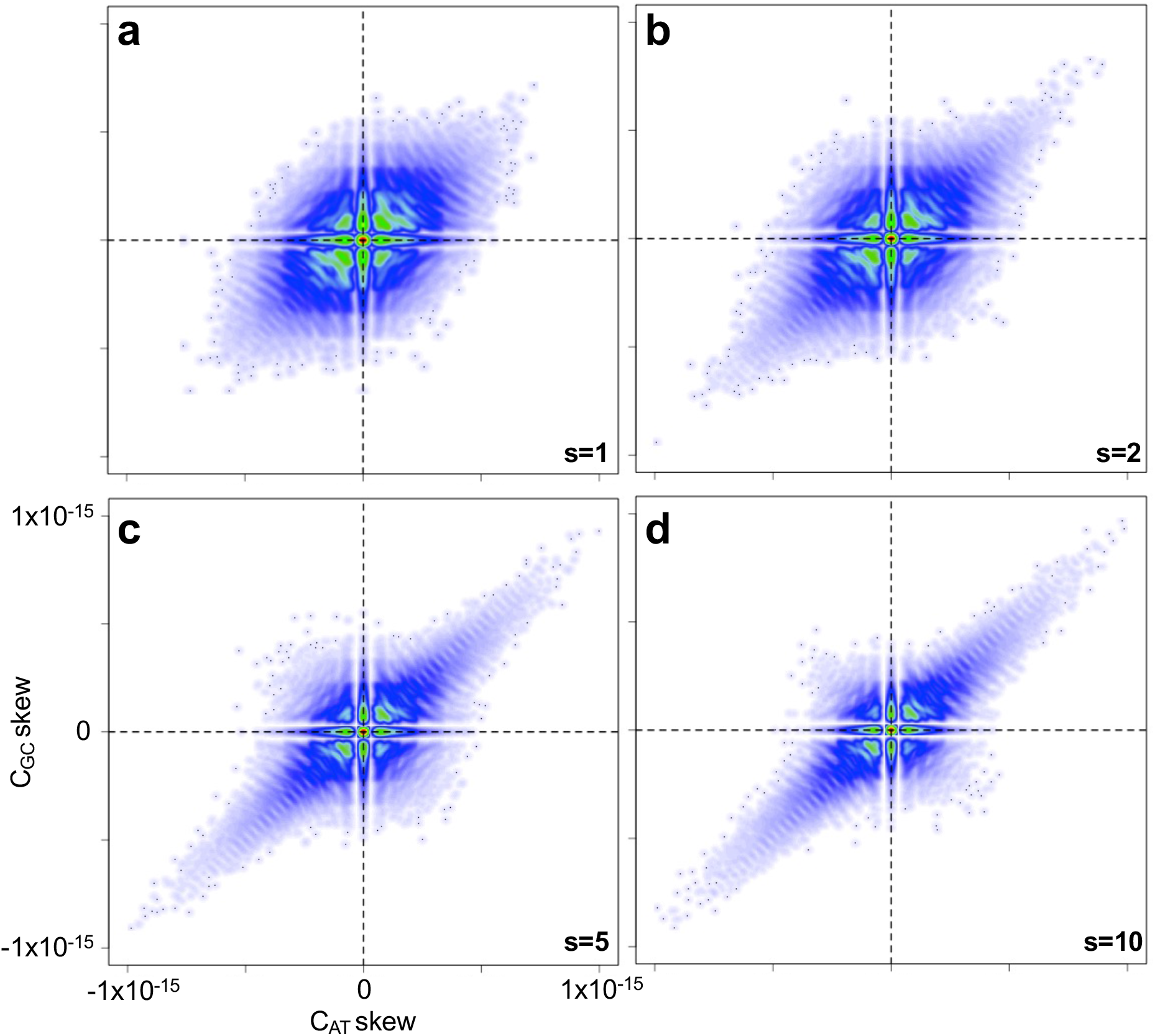
Numerical analysis of the NSB model symmetry-constrained mutation rate constants assumed to have a normal distribution. The systems of equations were solved starting from 25% initial contents for all four bases and symmetry-constrained rate constants randomly drawn from a normal distribution obtained from the Trek methodology [16] in byr^−1^ range. 25,000,000 such systems were calculated to produce genomic base contents at the final 4.28-byr time point. The 2-dimensional kernel density estimate scatterplot in c presents the distribution of C_GC_ and C_AT_ content skews from the outcome of the simulation (colours vary with decreasing occurrence frequency from red to blue), with the white dotted box indicating the 1.5x zone of compliance for the DNA virus kingdom with PR-2. The simulations were run four times separately with standard deviation values additionally scaled by m ∈ {1,2,5,10} multipliers in plots a-d, respectively.

**Figure S5.**
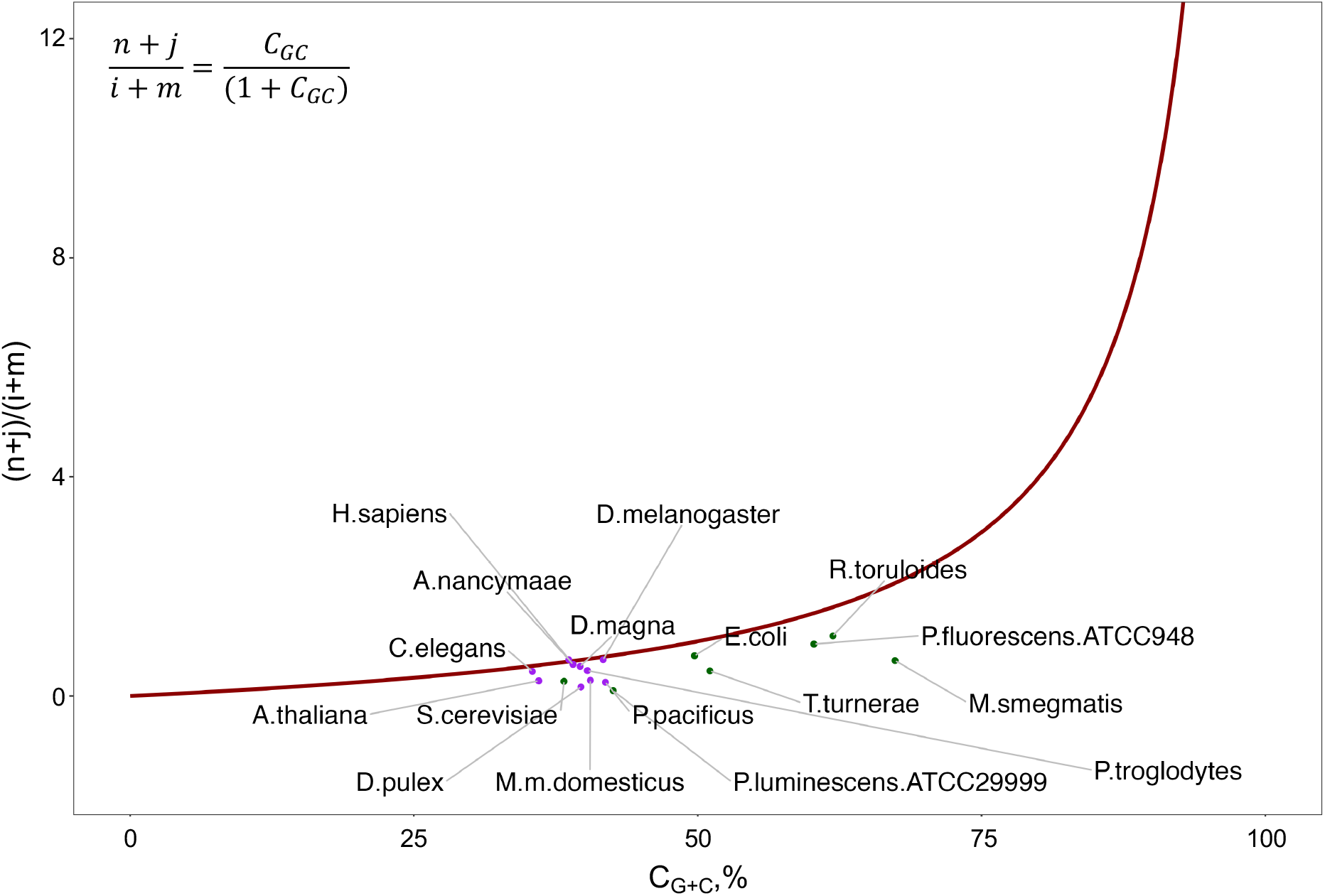
The base content solutions at equilibrium reveal the dependencies between the genomic C_G+C_ content and the mutation rate constants (red line). The strand symmetric mutation rate constants were obtained from 17 species across the eukaryotic and prokaryotic kingdoms [20–30] and overlaid as a scatterplot with colourings based on their associated eukaryotic (purple) or prokaryotic (dark green) kingdoms.

**Figure S6.**
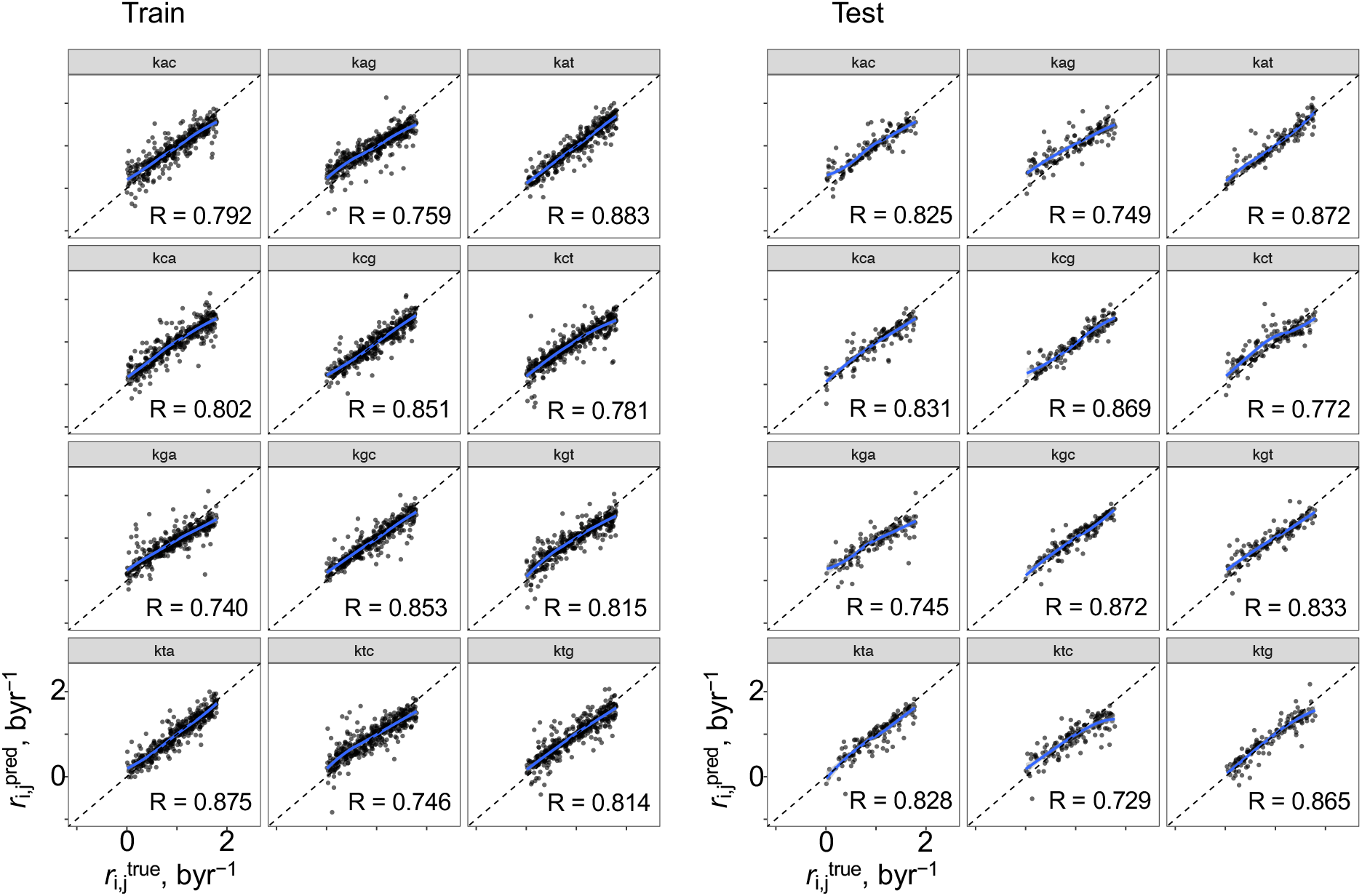
Evaluations of the analytical relations of the 12 mutation rate constants compliant with PR-2. (**left**) Equations generated from Eureqa (DataRobot) [31, 32] used to predict the mutation rate constant and comparing it against the mutation rate constant that arrived to the PR-2 compliant solution in the simulation. Pearson correlation coefficients are indicated on each graph where the equations for the k_A→T_ has the highest value (R = 0.883) while k_G→A_ has the lowest (R = 0.740). (**right**) Similar graphs were obtained for the test set (see main paper), where the equations for the k_G→C_ has the highest value (R = 0.872) (R = 0.872 for k_A→T_) while k_T→C_ has the lowest (R = 0.729). The diagonal dotted line represents perfect correlation with a slope of 1.

**Figure S7.**
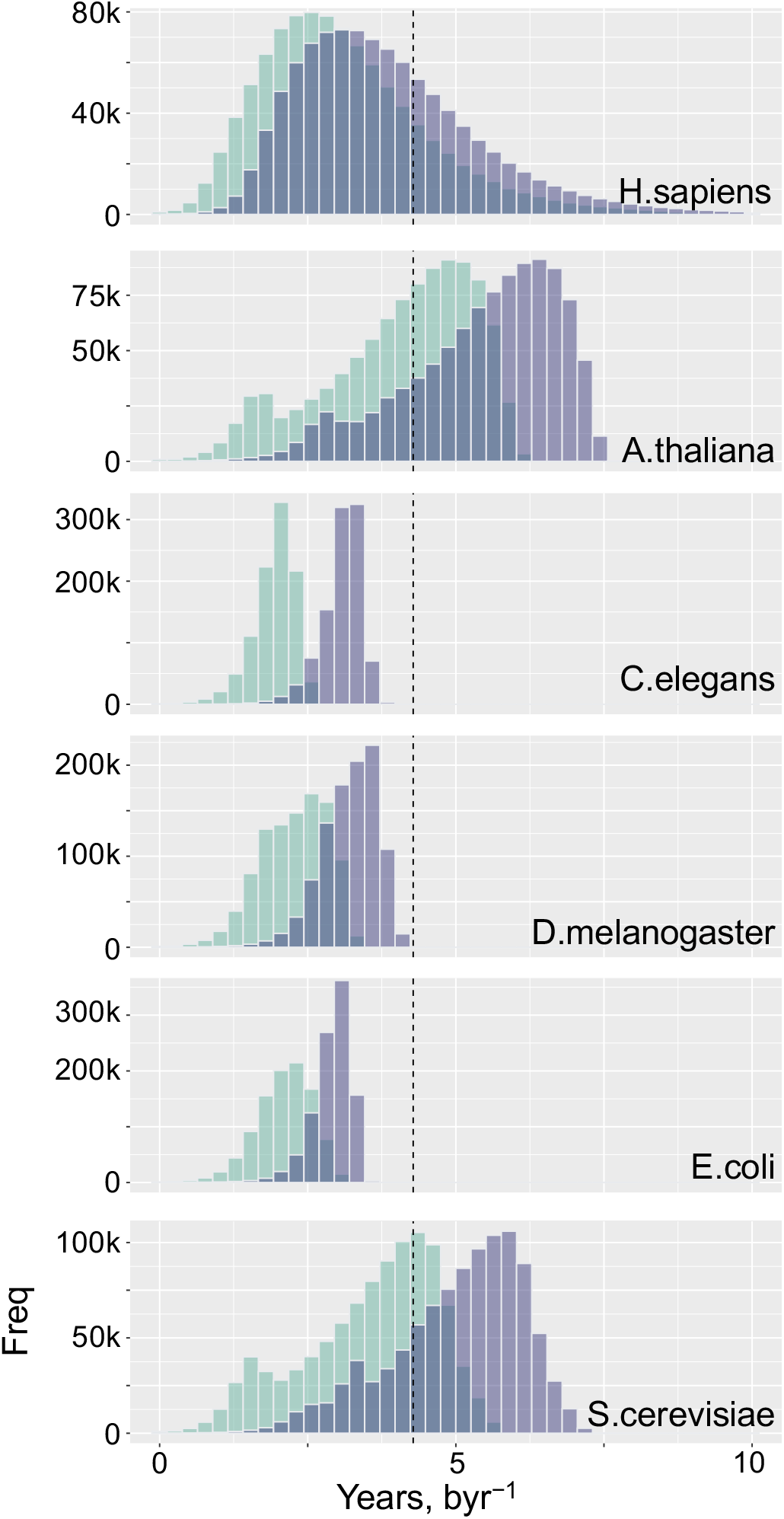
Distribution of time to reach PR-2 compliance and genome equilibrium. 10 million systems with the simulation model were generated where the initial base content was randomly sampled from the maximum allowed range based on prokaryotic organisms and the strand symmetric-based mutation rate constants were randomly drawn from a truncated normal distribution, the mutation rate constants of which were obtained from work done by Michael Lynch [30]. This process was performed for each of the six species independently. The green histograms represent the distribution of time to reach PR-2 compliance and the purple histograms represent the distribution of time to reach genome equilibrium (see **Materials and Methods** for details). The vertical dotted line intercepts the x-axis at 4.28 billion years, the maximum current estimate of age of life on Earth [33].

**Figure S8.**
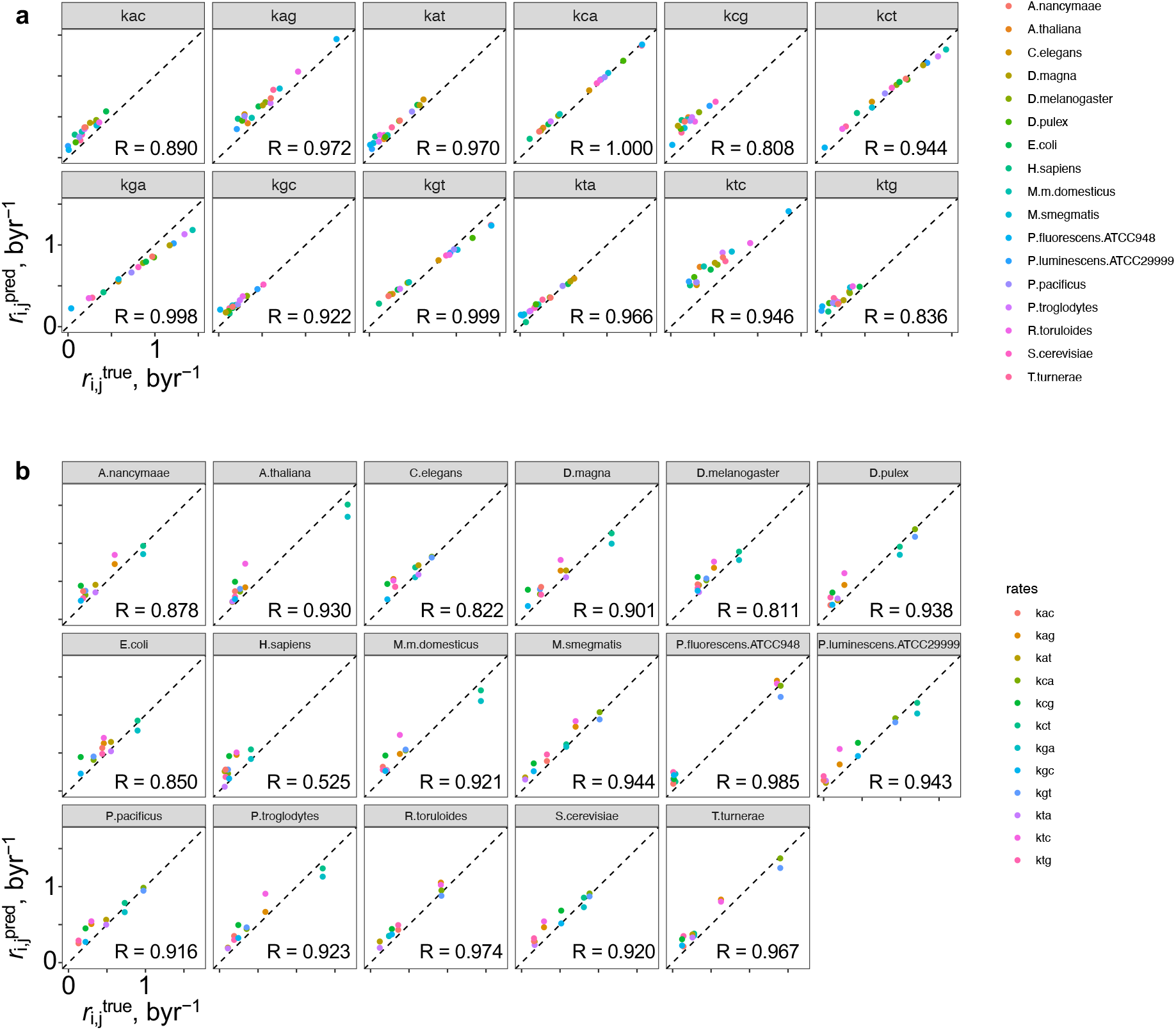
Examination of the closeness of current PR-2 compliant lifeforms to the fully equilibrated solution. (**a**) The strand symmetric mutation rate constants were obtained from 17 species across the eukaryotic and prokaryotic kingdoms [20–30], aligned to the Trek-scaling [16] and substituted the parameters of the 12 sets of mutation rate constant equations. Each species is represents a unique colour. The Pearson correlation coefficient values are indicated on each graph where the k_C→A_ obtained the highest value (R = 1.000). (**b**) Similar graphs were obtained where each graph represents all the 12 mutation rate constants from (**a**) for each of the 17 species. The Homo sapiens species has the lowest Pearson correlation coefficient (R = 0.525). The diagonal dotted line represents perfect correlation with a slope of 1.

## REFERENCES

1. Chargaff E. Chemical specificity of nucleic acids and mechanism of their enzymatic degradation. Experientia. 1950 Jun;6(6):201–9.

2. Watson JD, Crick FHC. Molecular structure of nucleic acids: A structure for deoxyribose nucleic acid. Nature. 1953;171(4356):737–8.

3. Rudner R, Karkas JD, Chargaff E. Separation of B. subtilis DNA into complementary strands, I. Biological properties. Proc Natl Acad Sci. 1968 Jun;60(2):630–5.

4. Rudner R, Karkas JD, Chargaff E. Separation of B. subtilis DNA into complementary strands. 3. Direct analysis. Proc Natl Acad Sci. 1968;60(3):921–2.

5. Prabhu VV. Symmetry observations in long nucleotide sequences. Nucleic Acids Res. 1993 Jun;21(12):2797–2797.

6. Lobry JR. Properties of a general model of DNA evolution under no-strand-bias conditions. J Mol Evol. 1995;40:326–30.

7. Baisnée PF, Hampson S, Baldi P. Why are complementary DNA strands symmetric? Bioinformatics. 2002 Aug;18(8):1021–33.

8. Shporer S, Chor B, Rosset S, Horn D. Inversion symmetry of DNA k-mer counts: Validity and deviations. BMC Genomics. 2016 Aug;17(1):696–696.

9. Fariselli P, Taccioli C, Pagani L, Maritan A. DNA sequence symmetries from randomness: the origin of the Chargaff’s second parity rule. Brief Bioinform. 2020 Apr;2020(00):1–10.

10. Cristadoro G, Degli Esposti M, Altmann EG. The common origin of symmetry and structure in genetic sequences. Sci Rep. 2018;8:15817–15817.

11. Nikolaou C, Almirantis Y. Deviations from Chargaff’s second parity rule in organellar DNA. Insights into the evolution of organellar genomes. Gene. 2006 Oct;381(1–2):34–41.

12. Mitchell D, Bridge R. A test of Chargaff’s second rule. Biochem Biophys Res Commun. 2006 Feb;340(1):90–4.

13. Okamura K, Wei J, Scherer SW. Evolutionary implications of inversions that have caused intra-strand parity in DNA. BMC Genomics. 2007 Jun;8(1):1–7.

14. Sueoka N. Intrastrand parity rules of DNA base composition and usage biases of synonymous codons. J Mol Evol. 1995;40:318–25.

15. Forsdyke DR. A stem-loop ‘kissing’ model for the initiation of recombination and the origin of introns. Mol Biol Evol. 1995;12(5):949–58.

16. Forsdyke DR. Relative roles of primary sequence and (G + C)% in determining the hierarchy of frequencies of complementary trinucleotide pairs in DNAs of different species. J Mol Evol. 1995;41:573–81.

17. Hart A, Servet Martínez ·, Olmos F. A Gibbs approach to Chargaff’s second parity rule. J Stat Phys. 2012;146:408–22.

18. Afreixo V, Rodrigues JMOS, Bastos CAC. Analysis of single-strand exceptional word symmetry in the human genome: New measures. Biostatistics. 2015 Jul;16(2):209–21.

19. Afreixo V, Rodrigues JMOS, Bastos CAC, Tavares AHMP, Silva RM. Exceptional symmetry by genomic word: A statistical analysis. Interdiscip Sci Comput Life Sci. 2017 Mar;9(1):14–23.

20. Albrecht-Buehler G. Inversions and inverted transpositions as the basis for an almost universal ‘format’ of genome sequences. Genomics. 2007 Sep;90(3):297–305.

21. Zhang SH, Huang YZ. Characteristics of oligonucleotide frequencies across genomes: Conservation versus variation, strand symmetry, and evolutionary implications. Nat Preced 2008. 2008 Aug;1–1.

22. R Core Team. R: A language and environment for statistical computing [Internet]. Vienna, Austria: R Foundation for Statistical Computing; 2021. Available from: https://www.R-project.org/

23. Chang W, Cheng J, Allaire J, Sievert C, Schloerke B, Xie Y, et al. shiny: Web application framework for R. R package version 1.7.1 [Internet]. 2021. Available from: https://CRAN.R-project.org/package=shiny

24. Wickham H. ggplot2: Elegant graphics for data analysis [Internet]. Springer-Verlag New York; 2016. Available from: https://ggplot2.tidyverse.org

25. Baptiste A. gridExtra: Miscellaneous functions for ‘grid’ graphics [Internet]. 2017. Available from: https://CRAN.R-project.org/package=gridExtra

26. Wickham H, Averick M, Bryan J, Chang W, McGowan LD, François R, et al. Welcome to the tidyverse. J Open Source Softw. 2019 Nov 21;4(43):1686.

27. Wickham H. Reshaping data with the reshape package. J Stat Softw. 2007 Nov 13;21:1–20.

28. Stabenau A, McVicker G, Melsopp C, Proctor G, Clamp M, Birney E. The ensembl core software libraries. Genome Res. 2004 Jan 5;14(5):929–33.

29. Sahakyan AB, Balasubramanian S. Single genome retrieval of context-dependent variability in mutation rates for human germline. BMC Genomics [Internet]. 2017 Jan;18(1). Available from: /pmc/articles/PMC5237266/

30. Lynch M. Rate, molecular spectrum, and consequences of human mutation. Proc Natl Acad Sci. 2010 Jan;107(3):961–8.

31. Mersmann O, Trautmann H, Steuer D, Bornkamp B. truncnorm: Truncated normal distribution [Internet]. 2018. Available from: https://CRAN.R-project.org/package=truncnorm

32. Microsoft, Weston S. foreach: Provides foreach looping construct [Internet]. 2020. Available from: https://CRAN.R-project.org/package=foreach

33. Microsoft Corporation, Weston S. doParallel: Foreach parallel adaptor for the ‘parallel’ package [Internet]. 2020. Available from: https://CRAN.R-project.org/package=doParallel

34. Gaujoux R. doRNG: Generic reproducible parallel backend for ‘foreach’ loops [Internet]. 2020. Available from: https://CRAN.R-project.org/package=doRNG

35. Sung W, Ackerman MS, Gout JF, Miller SF, Williams E, Foster PL, et al. Asymmetric context-dependent mutation patterns revealed through mutation-accumulation experiments. Mol Biol Evol. 2015 Jul 1;32(7):1672–83.

36. Lobry JR, Lobry C. Evolution of DNA base composition under no-strand-bias conditions when the substitution rates are not constant. Mol Biol Evol. 1999 Jan 1;16(6):719–23.

37. Lobry JR. A nice wrong model for the evolution of DNA base frequencies. Phys A. 1999;273:99–102.

38. Reijns MAM, Kemp H, Ding J, Marion de Procé S, Jackson AP, Taylor MS. Lagging-strand replication shapes the mutational landscape of the genome. Nature. 2015 Feb;518(7540):502–6.

39. Zhang SH, Huang YZ. Limited contribution of stem-loop potential to symmetry of single-stranded genomic DNA. Bioinformatics. 2009 Dec;26(4):478–85.

40. Chen L, Zhao H. Negative correlation between compositional symmetries and local recombination rates. Bioinformatics. 2005 Nov;21(21):3951–8.

41. Zhang SH, Wang L. A novel common triplet profile for GC-rich prokaryotic genomes. Genomics. 2011 May;97(5):330–1.

42. Agier N, Fischer G. The mutational profile of the yeast genome is shaped by replication. Mol Biol Evol. 2012 Mar 1;29(3):905–13.

43. Dodd MS, Papineau D, Grenne T, Slack JF, Rittner M, Pirajno F, et al. Evidence for early life in Earth’s oldest hydrothermal vent precipitates. Nature. 2017 Mar;543(7643):60–4.

44. Senra MVX, Sung W, Ackerman M, Miller SF, Lynch M, Soares CAG. An unbiased genome-wide view of the mutation rate and spectrum of the Endosymbiotic Bacterium Teredinibacter turnerae. Genome Biol Evol. 2018 Mar 1;10(3):723–30.

45. Ho EKH, Macrae F, Latta LC 4th, McIlroy P, Ebert D, Fields PD, et al. High and highly variable spontaneous mutation rates in Daphnia. Mol Biol Evol. 2020 Nov 1;37(11):3258–66.

46. Long H, Sung W, Miller SF, Ackerman MS, Doak TG, Lynch M. Mutation rate, spectrum, topology, and context-dependency in the DNA mismatch repair-deficient Pseudomonas fluorescens ATCC948. Genome Biol Evol. 2015 Jan 1;7(1):262–71.

47. Weller AM, Rödelsperger C, Eberhardt G, Molnar RI, Sommer RJ. Opposing forces of A/T-biased mutations and G/C-biased gene conversions shape the genome of the Nematode Pristionchus pacificus. Genetics. 2014 Apr 1;196(4):1145–52.

48. Thomas GWC, Wang RJ, Puri A, Harris RA, Raveendran M, Hughes DST, et al. Reproductive longevity predicts mutation rates in Primates. Curr Biol. 2018 Oct 8;28(19):3193–3197.e5.

49. Dumont BL. Significant strain variation in the mutation spectra of inbred laboratory Mice. Mol Biol Evol. 2019 May 1;36(5):865–74.

50. Long H, Behringer MG, Williams E, Te R, Lynch M. Similar mutation rates but highly diverse mutation spectra in Ascomycete and Basidiomycete Yeasts. Genome Biol Evol. 2016 Dec 1;8(12):3815–21.

51. Flynn JM, Chain FJJ, Schoen DJ, Cristescu ME. Spontaneous mutation accumulation in Daphnia pulex in selection-free vs. competitive environments. Mol Biol Evol. 2017 Jan 1;34(1):160–73.

52. Pan J, Williams E, Sung W, Lynch M, Long H. The insect-killing bacterium Photorhabdus luminescens has the lowest mutation rate among bacteria. Mar Life Sci Technol. 2021 Feb 1;3(1):20–7.

53. Kucukyildirim S, Long H, Sung W, Miller SF, Doak TG, Lynch M. The rate and spectrum of spontaneous mutations in Mycobacterium smegmatis, a Bacterium naturally devoid of the postreplicative mismatch repair pathway. G3 GenesGenomesGenetics. 2016 Jul 1;6(7):2157–63.

54. Schmidt MD, Lipson H. Coevolution of fitness predictors. IEEE Trans Evol Comput. 2008;12(6).

55. Schmidt M, Lipson H. Distilling free-form natural laws from experimental data. Science. 2009 Apr;324(5923):81–5.

## References

[1] David N. Cooper et al. “The CpG dinucleotide and human genetic disease”. In: Human Genetics 78.2 (Feb. 1988). Publisher: Springer, pp. 151–155. doi: 10.1007/BF00278187.

[2] Adrian P. Bird. “DNA methylation and the frequency of CpG in animal DNA”. In: Nucleic Acids Research 8.7 (Apr. 1980). Publisher: Oxford Academic, pp. 1499–1504. doi: 10.1093/NAR/8.7.1499.

[3] Christine Coulondre et al. “Molecular basis of base substitution hotspots in Escherichia coli”. In: Nature 274.5673 (1978). Publisher: Nature Publishing Group, pp. 775–780. doi: 10.1038/274775a0.

[4] Michael W. Nachman et al. “Estimate of the mutation rate per nucleotide in Humans”. In: Genetics 156.1 (Sept. 2000), pp. 297–304. doi: 10.1093/genetics/156.1.297.

[5] Cizhong Jiang et al. “Directionality of point mutation and 5-methylcytosine deamination rates in the chimpanzee genome”. In: BMC Genomics 7.1 (Dec. 2006). Publisher: BioMed Central, pp. 1–13. doi: 10.1186/1471-2164-7-316.

[6] Zhao Z et al. “Worldwide DNA sequence variation in a 10-kilobase noncoding region on human chromosome 22”. In: Proceedings of the National Academy of Sciences of the United States of America 97.21 (Oct. 2000). Publisher: Proc Natl Acad Sci U S A, pp. 11354–11358. doi: 10.1073/PNAS.200348197.

[7] Feng-Chi Chen et al. “Genomic divergences between Humans and other Hominoids and the effective population size of the common ancestor of Humans and Chimpanzees”. In: American Journal of Human Genetics 68.2 (2001). Publisher: Elsevier, pp. 444–444. doi: 10.1086/318206.

[8] Jerome H. Friedman. “Greedy function approximation: A gradient boosting machine.” In: Institute of Mathematical Statistics 29.5 (Oct. 2001). Publisher: Institute of Mathematical Statistics, pp. 1189–1232. doi: 10.1214/AOS/1013203451.

[9] Alexey Natekin et al. “Gradient boosting machines, a tutorial”. In: Frontiers in Neurorobotics 7.DEC (2013). Publisher: Frontiers Media SA. doi: 10.3389/FNBOT.2013.00021.

[10] Baoshan Ma et al. “Diagnostic classification of cancers using extreme gradient boosting algorithm and multi-omics data”. In: Computers in Biology and Medicine 121 (June 2020). Publisher: Pergamon, pp. 103761–103761. doi: 10.1016/J.COMPBIOMED.2020.103761.

[11] Max Kuhn. “Building predictive models in R using the caret package”. In: Journal of Statistical Software 28 (Nov. 2008), pp. 1–26. doi: 10.18637/jss.v028.i05.

[12] Christopher John R. MLeval: Machine learning model evaluation. 2020.

[13] Jesse Davis et al. “The relationship between precision-recall and ROC curves”. In: ACM International Conference Proceeding Series 148 (2006), pp. 233–240. doi: 10.1145/1143844.1143874.

[14] László A. Jeni et al. “Facing imbalanced data - recommendations for the use of performance metrics”. In: Proceedings - 2013 Humaine Association Conference on Affective Computing and Intelligent Interaction, ACII 2013 (2013), pp. 245–251. doi: 10.1109/ACII.2013.47.

[15] Bowen Song et al. “ROC operating point selection for classification of imbalanced data with application to computer-aided polyp detection in CT colonography”. In: International journal of computer assisted radiology and surgery 9.1 (Jan. 2014). Publisher: NIH Public Access, pp. 79–79. doi: 10.1007/S11548-013-0913-8.

[16] Aleksandr B. Sahakyan et al. “Single genome retrieval of context-dependent variability in mutation rates for human germline”. In: BMC Genomics 18.1 (Jan. 2017). Publisher: BioMed Central. doi: 10.1186/S12864-016-3440-5.

[17] Liangliang Liu et al. “An interpretable boosting model to predict side effects of analgesics for osteoarthritis”. In: BMC Systems Biology 12.6 (Nov. 2018). Publisher: BioMed Central, pp. 29–38. doi: 10.1186/S12918-018-0624-4.

[18] Aleksandr B. Sahakyan et al. “Machine learning model for sequence-driven DNA G-quadruplex formation”. In: Scientific Reports 7.1 (Nov. 2017). Publisher: Nature Publishing Group, pp. 1–11. doi: 10.1038/s41598-017-14017-4.

[19] Bruce G. Marcot et al. “What is an optimal value of k in k-fold cross-validation in discrete Bayesian network analysis?” In: Computational Statistics 36.3 (June 2020). Publisher: Springer, pp. 2009–2031. doi: 10.1007/S00180-020-00999-9.

[20] Jiao Pan et al. “The insect-killing bacterium Photorhabdus luminescens has the lowest mutation rate among bacteria”. In: Mar Life Sci Technol 3.1 (Feb. 2021), pp. 20–27. doi: 10.1007/s42995-020-00060-0.

[21] Beth L Dumont. “Significant strain variation in the mutation spectra of inbred laboratory Mice”. In: Molecular Biology and Evolution 36.5 (May 2019), pp. 865–874. doi: 10.1093/molbev/msz026.

[22] Eddie K H Ho et al. “High and highly variable spontaneous mutation rates in Daphnia”. In: Molecular Biology and Evolution 37.11 (Nov. 2020), pp. 3258–3266. doi: 10.1093/molbev/msaa142.

[23] Gregg W. C. Thomas et al. “Reproductive longevity predicts mutation rates in Primates”. In: Current Biology 28.19 (Oct. 2018), 3193–3197.e5. doi: 10.1016/j.cub.2018.08.050.

[24] Marcus V X Senra et al. “An unbiased genome-wide view of the mutation rate and spectrum of the Endosymbiotic Bacterium Teredinibacter turnerae”. In: Genome Biology and Evolution 10.3 (Mar. 2018), pp. 723–730. doi: 10.1093/gbe/evy027.

[25] Jullien M. Flynn et al. “Spontaneous mutation accumulation in Daphnia pulex in selection-free vs. competitive environments”. In: Molecular Biology and Evolution 34.1 (Jan. 2017), pp. 160–173. doi: 10.1093/molbev/msw234.

[26] Hongan Long et al. “Similar mutation rates but highly diverse mutation spectra in Ascomycete and Basidiomycete Yeasts”. In: Genome Biology and Evolution 8.12 (Dec. 2016), pp. 3815–3821. doi: 10.1093/gbe/evw286.

[27] Sibel Kucukyildirim et al. “The rate and spectrum of spontaneous mutations in Mycobacterium smegmatis, a Bacterium naturally devoid of the postreplicative mismatch repair pathway”. In: G3 Genes—Genomes—Genetics 6.7 (July 2016), pp. 2157–2163. doi: 10.1534/g3.116.030130.

[28] Hongan Long et al. “Mutation rate, spectrum, topology, and context-dependency in the DNA mismatch repair-deficient Pseudomonas fluorescens ATCC948”. In: Genome Biology and Evolution 7.1 (Jan. 2015), pp. 262–271. doi: 10.1093/gbe/evu284.

[29] Andreas M Weller et al. “Opposing forces of A/T-biased mutations and G/C-biased gene conversions shape the genome of the Nematode Pristionchus pacificus”. In: Genetics 196.4 (Apr. 2014), pp. 1145–1152. doi: 10.1534/genetics.113.159863.

[30] Michael Lynch. “Rate, molecular spectrum, and consequences of human mutation”. In: PNAS 107.3 (Jan. 2010). Publisher: National Academy of Sciences Section: Biological Sciences, pp. 961–968. doi: 10.1073/pnas.0912629107.

[31] Michael D Schmidt et al. “Coevolution of fitness predictors”. In: IEEE Transactions on Evolutionary Computation 12.6 (2008). doi: 10.1109/TEVC.2008.919006.

[32] Michael Schmidt et al. “Distilling free-form natural laws from experimental data”. In: Science 324.5923 (Apr. 2009), pp. 81–85. doi: 10.1126/SCIENCE.1165893.

[33] Matthew S. Dodd et al. “Evidence for early life in Earth’s oldest hydrothermal vent precipitates”. In: Nature 543.7643 (Mar. 2017), pp. 60–64. doi: 10.1038/nature21377.

